# C-terminal domain of the filamentous hemagglutinin FhaB is crucial for interaction of *Bordetella pertussis* with ciliated epithelial cells

**DOI:** 10.64898/2026.05.25.727647

**Authors:** David Jurnecka, Josef Chmelik, Jana Holubova, Ondrej Stanek, Petra Kasparova, Abdul Samad, Jana Proskova, Karolina Skopova, Marketa Dalecka, Dominik Pinkas, Peter Sebo, Christopher S. Hayes, Ladislav Bumba

## Abstract

*Bordetella pertussis*, the causative agent of whooping cough, produces a ∼370 kDa filamentous hemagglutinin FhaB that serves as a major bacterial adhesin in airway infection. FhaB is secreted via a two-partner secretion pathway and under *in vitro* culture conditions it is proteolytically processed to the shed ∼230 kDa FHA antigen used currently in acellular pertussis vaccines. We show that FhaB remains largely unprocessed during *B. pertussis* adhesion to ciliated airway epithelial cells and that its C-terminal domain (CT) is essential for the adhesin function of FhaB. CT deletion did not affect FhaB folding, secretion, or surface exposure, but abolished *B. pertussis* adhesion to primary human nasal ciliated epithelial cells, thus preventing bacterial colonization of the nasal mucosa and shedding and transmission of the pathogen in a murine nasal infection model. *In situ* cryo–electron tomography revealed a structural reorganization of the FhaB filaments upon contact with the cilia, presumably due to export of the CT from bacterial periplasm and its subsequent delivery across the ciliary membrane. These findings establish the CT of FhaB as a critical determinant of upper airway colonization by *B. pertussis* and identify the unprocessed FhaB as the biologically relevant adhesin form involved in airway infection. The revised model of FhaB biogenesis underpins its unique mode of action in pertussis pathogenesis and makes the CT domain to a candidate antigen for future pertussis vaccines.

## Introduction

*Bordetella pertussis* is a Gram-negative coccobacillus that elicits a highly contagious respiratory illness known as pertussis, or whooping cough (1–3). Before the introduction of generalized vaccination, pertussis used to be the leading cause of infant mortality due to infectious diseases and accounted for more infant deaths than all the other infections combined (4, 5). Despite global immunization programs, pertussis remains among the least-controlled vaccine-preventable diseases, with dozens of millions of whooping cough cases and hundreds of thousands of pertussis-related deaths occurring annually worldwide (6, 7). *B. pertussis* produces an extensive repertoire of virulence factors, including immunomodulatory protein toxins like pertussis toxin, adenylate cyclase toxin, and dermonecrotic toxin, as well as factors involved in complement resistance, iron acquisition, and adhesion to host epithelial and immune cells (8–13). The collective action of these virulence determinants results in suppression of the innate and modulation of the adaptive immune responses. The ensuant immune evasion promotes long-term colonization of the upper airway mucosa and enables *B. pertussis* to circulate in highly vaccinated populations (14). A prominent role in this capacity is played by the adhesive fimbriae and filamentous hemagglutinin (FhaB) appendages that act in concert to mediate a tight and highly selective attachment of the bacteria to the motile cilia of airway epithelial cells (15–18).

FhaB is transported across the bacterial outer membrane by one of the two-partner secretion (TPS) systems that excrete large proteins with diverse biological functions (19, 20). The *B. pertussis fhaB* gene encodes a precursor protein of 3590 amino acid residues with a 71 residue-long N-terminal signal peptide that directs the Sec-dependent translocation of preFhaB into the periplasm (Fig. 1A)(21, 22). Upon signal peptide cleavage, the conserved N-terminal TPS domain of FhaB engages the polypeptide transport-associated (POTRA) domains of its cognate outer membrane transporter FhaC (23–25). An about ∼2400 residue-long FhaB polypeptide segment then loops out through the FhaC pore, forming a hairpin-like structure of an elongated N- to C- folded β stack. This is capped by a globular structure previously called the ‘mature C-terminal domain’ (MCD), which is connected by a latch across the FhaC pore to a C-terminal FhaB moiety of ∼1240 residues. For historical reasons, this was erroneously called a ‘prodomain’ (26, 27), which was proposed to be retained in the periplasmic space by its ‘prodomain N-terminal’ (PNT) segment and eventually undergo proteolytic degradation by a complex sequence of processing steps (28). This is thought to be initiated by cleavage of the C-terminal (CT) domain by the periplasmic protease DegP, followed by progressive degradation of the rest of the ‘prodomain’ by the carboxy-terminal protease CtpA (29, 30). Degradation of the C-terminal ‘prodomain’ would then enable processing of the protruding FhaB hairpin by the subtilisin-like autotransporter protease SphB1 to the ∼230 kDa N-terminal ‘mature’ FHA fragment shed into extracellular milieu (31, 32). However, the biological role of SphB1-mediated processing of FhaB remains unclear and other bacterial proteases of SphB1-deficient *Bordetella* can generate reduced levels of a slightly larger ∼250 kDa FHA fragment (33).

**Figure 1.**
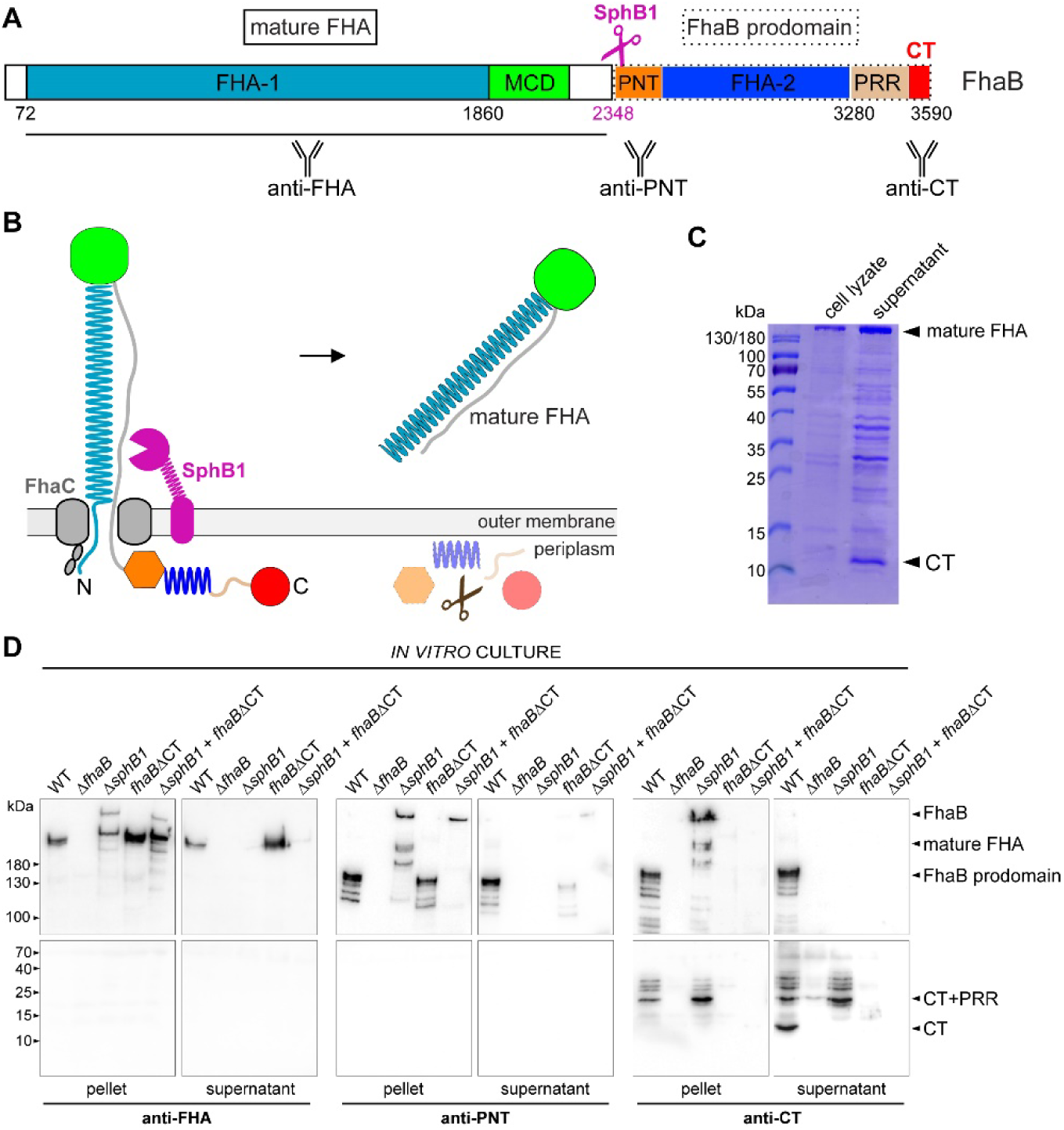
The C-terminal FhaB ‘prodomain’ moiety is not completely degraded in the periplasm. (A) Schematic representation of FhaB domain architecture. FHA-1 and FHA-2 denote regions containing tandem 19–amino acid repeats predicted to form a right-handed β-helical shaft. MCD, mature C-terminal domain; PNT, ‘prodomain’ N terminus; CT, C-terminal domain. The epitope map of polyclonal sera is indicated. (B) Current model of FhaB biogenesis and maturation. Translocation of FhaB across the outer membrane is initiated by interaction of the N-terminal TPS domain with the periplasmic POTRA domains of the cognate transporter FhaC. As FhaB loops out from the bacterium in an N- to C-terminal hairpin configuration, tandem repeats within the FHA-1 domain sequentially fold into a β-helical shaft, forming an elongated filament capped by the globular mature C-terminal domain (MCD). Following cleavage of FhaB by the surface-anchored protease SphB1, mature FHA is released, whereas the FhaB ‘prodomain’ is retained in the periplasm and is thought to be rapidly degraded. (C) SDS–PAGE analysis of cell lysate and the concentrated culture supernatants from overnight cultures of *B. pertussis*. Bands corresponding to mature FHA and CT are indicated. (D) Immunoblot analysis of cell lysates and culture supernatants from overnight cultures of the indicated *B. pertussis* strains grown in SS medium at 37 °C. The lysates were separated on 5 % (upper panels) and 15 % (lower panels) SDS-PAGE gels and membranes containing transferred proteins were probed with anti-FHA, anti-PNT, and anti-CT polyclonal mouse sera. Bands corresponding to distinct FhaB forms are indicated.

The released FHA fragment is an elongated and rigid rod-shaped molecule of approximately 50 nm in length and 4 nm in diameter (34). Structural analysis of the N-terminal TPS domain of FHA revealed that its tandem 19 residue-long repeats are folded into a right-handed parallel β-helical coil that is composed of three parallel β-sheets arranged in a triangular cross-section (35). This β-helix architecture is predicted to extend along the FHA molecule up to its C-terminal MCD cap (36). Four distinct functional domains of the FHA fragment were identified and proposed to cooperate in FhaB-mediated *Bordetella* adhesion to various host cells (37, 38). A heparin-binding domain, comprising residues 440-860 appears to account for the hemagglutinating activity of FHA and would facilitate FhaB-mediated adhesion to sulfated glycosaminoglycans of extracellular matrix and host cell surface glycoproteins (39–41). A conserved Arg-Gly-Asp (RGD) motif in the FHA segment of FhaB (residues 1096-1098) was proposed to participate in *B. pertussis* adhesion to macrophages (42). The carbohydrate recognition domain (residues 1140-1280) has been implicated in interactions with galactose-containing glycoconjugates and galactose-glucose-containing glycolipids (43, 44). Finally, the MCD tip was implicated in *Bordetella* adhesion to macrophage-like and epithelial cells and was reported to be required for colonization of the upper respiratory tract of rodents (28, 45, 46). In addition to its role in bacterial adhesion, FhaB and its FHA fragment have also been reported to play a role in modulation of host immune responses, inducing both pro- and anti-inflammatory cytokines and contributing to suppression of mucosal inflammation during bacterial colonization (47–52). However, the molecular basis of FhaB/FHA interactions with cellular receptors and the ensuing signaling remain unclear and a matter of debate (17, 45, 53).

Curiously, the released FHA fragment has historically been regarded as the primary functional form of *Bordetella* filamentous hemagglutinin, even though its role as an adhesin was inferred from cell-binding experiments with *Bordetella* bacteria producing full-length FhaB. This conceptual conundrum of a shed molecule playing a major role in bacterial adhesion has recently been resolved by the evidence that full-length FhaB plays an essential role during airway infection. Deletion of the ∼100 residue-long C-terminal CT segment, or of the adjacent proline-rich region of ‘prodomain’, strongly impaired the capacity of *B. bronchiseptica* mutants to persist in the lower respiratory tract of mice (54). Moreover, the primary sequence of CT is fully conserved in airway-infecting *Bordetella* species and this domain was recently found to bind microtubules and play a crucial role in the movement of *B. bronchiseptica* bacteria along the beating cilia of airway epithelial cells (55). Here, we visualize the structure of cell-bound FhaB by cryo-EM tomography, present the atomic structure of CT solved by nuclear magnetic resonance spectroscopy and demonstrate that the CT domain is essential for bacterial binding to human ciliated nasal epithelial cells and for nasal cavity colonization by *B. pertussis*.

## Results

### The C-terminal moiety of FhaB is translocated and released into the extracellular milieu

The current model of FhaB biogenesis proposes that about two thirds of the FhaB polypeptide are translocated to the bacterial surface through the FhaC pore. Upon processing by SphB1, a shed N-terminal ∼230-kDa FHA fragment is generated, while the C-terminal ‘prodomain’ moiety (∼130 kDa) is retained and degraded within the bacterial periplasm (Fig. 1B). However, full-length FhaB has recently been identified as the functional form required for airway colonization by *Bordetella* (54, 55). We thus reassessed the fate of the FhaB ‘prodomain’ by proteomic analysis of tryptic digests derived from cellular (pellet) and culture supernatant fractions of *B. pertussis* liquid cultures. In the cellular fraction, ∼56% sequence coverage spanning the entire FhaB polypeptide was achieved (Fig. S1), whereas in culture supernatants ∼66% coverage of the released ‘mature’ FHA (residues 72–2348) was detected (Fig. S2). Unexpectedly, the supernatant fraction also contained peptides covering ∼37% of the FhaB ‘prodomain’ sequence, particularly within its C-terminal segment (Fig. S2). These peptides originated from polypeptide species retained by a 30 kDa cut-off filter during sample preparation, and many were consecutive in sequence. This indicated that at least a part of the C-terminal moiety of FhaB was released into the extracellular milieu as a larger polypeptide that has not been degraded during transit through the periplasm. Accordingly, SDS–PAGE analysis of precipitated culture supernatants revealed several protein bands (Fig. 1C), with the most abundant ∼12-kDa species being unambiguously identified as the CT domain, yielding the tryptic peptides _3497_HVVQQQVQVLQR_3508_, _3532_LTDENGPKQTYTINR_3546_ and_3562_TTLGLEQTFR_3571_, respectively (Table 1). These results demonstrated that the CT constitutes a stable and protease-resistant protein that is released into the extracellular environment.

**Table 1.**
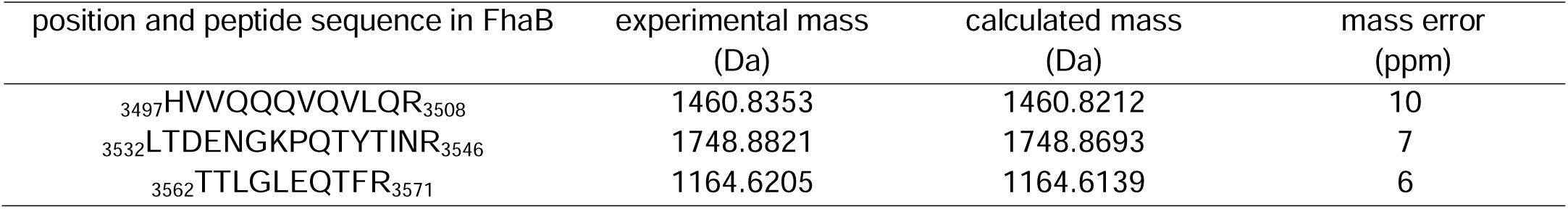
Peptide sequences obtained from the 12-kDa gel band.

To investigate the topology and fate of the FhaB ‘prodomain’ in more detail, we generated mouse polyclonal antisera against the PNT (residues 2349–2829 of FhaB) and CT (residues 3495–3590) segments. The relationship between FhaB ‘prodomain’ processing and mature FHA production was then examined by Western blot analysis of cellular and culture supernatant fractions using a panel of *B. pertussis* mutants. As shown in Fig. 1D, the *B. pertussis fhaB*ΔCT mutant, which expresses FhaB lacking the CT sequence, produced higher levels of mature FHA (∼230 kDa) associated with cells and released it into culture supernatants as the wild-type (*fhaB*^+^) strain. Thus, the absence of CT did not impair FHA processing but instead accelerated FhaB surface exposure and release of the FHA fragment. In contrast, deletion of the SphB1 protease gene (Δ*sphB1)* abolished the release of processed FHA into the culture supernatant and led to accumulation of full-length FhaB-protein (∼370 kDa) in the cellular fraction, together with a larger ∼250-kDa FhaB-derived fragment detectable with anti-FHA serum. These data indicate that in the absence of SphB1, full-length FhaB is retained and stabilized within the FhaC translocon and undergoes alternative processing by an unidentified protease that generates a ‘mature-like’ FHA fragment of ∼250 kDa. The same FhaB-derived protein species were also detected in cells of the Δ*sphB1*/*fhaB*ΔCT double mutant.

Probing with the anti-PNT serum confirmed its high specificity for the N-terminal region of the FhaB ‘prodomain’ (Fig. 1D, middle panels). This serum detected full-length FhaB (∼370 kDa) in the cellular fractions of both Δ*sphB1* and Δ*sphB1*/*fhaB*ΔCT mutants, but did not recognize the abundant mature FHA (∼230 kDa) present in the culture supernatants of the wild-type (*fhaB*^+^) and *fhaB*ΔCT strains. Notably, the supernatant of the Δ*sphB1*/*fhaB*ΔCT strain contained a detectable ∼370-kDa band corresponding to entire unprocessed FhaBΔCT protein. Lower amounts of this protein were also observed in the supernatant of the *fhaB*ΔCT strain, but not in that of the Δ*sphB1* strain, indicating that removal of CT accelerates FhaB secretion and facilitates release of unprocessed FhaB from the cells. Consistent with this, the cellular fraction of the Δ*sphB1* strain, but not that of the Δ*sphB1*/*fhaB*ΔCT mutant, contained multiple FhaB degradation products detected by the anti-PNT serum, suggesting that the presence of CT promotes proteolysis of FhaB retained within the FhaC translocon. Importantly, in lysates of the SphB1-producing *fhaB*^+^ and *fhaB*ΔCT strains, the anti-PNT serum detected a prominent ∼130-kDa fragment corresponding to the FhaB ‘prodomain’ generated during processing to FHA. Strikingly, this ∼130-kDa ‘prodomain’ fragment and its degradation products were also abundant in the culture supernatant of the wild-type strain, indicating that the entire FhaB polypeptide can be translocated across the FhaC pore and that its processing to mature FHA by SphB1 can occur in the extracellular environment. However, the stability and extracellular accumulation of the ‘prodomain’ depended on CT, as only low levels of the ΔCT ‘prodomain’ fragment were detected in the culture supernatant of the *fhaB*ΔCT strain.

Consistent with these findings, the anti-CT serum specifically recognized the C-terminal segment of the FhaB ‘prodomain’ and detected full-length FhaB (∼370 kDa) in cells of the Δ*sphB1* mutant, as well as the corresponding ∼130 kDa ‘prodomain’ fragment resulting from FhaB processing in cells and culture supernatants of the wild-type strain (Fig.1D, right panels). Unlike the anti-PNT serum, the anti-CT serum revealed a series of ‘prodomain’-derived degradation products in both cellular and supernatant fractions of wild-type and Δ*sphB1* strains. This suggested that proteolysis of the ‘prodomain’ released by SphB1-dependent processing (in wild type) or by an alternative protease (ΔsphB1) is initiated at the N terminus and proceeds progressively toward the C terminus of the ‘prodomain’. Notably, incomplete ‘prodomain’ degradation resulted in accumulation of multiple fragments ranging from 20 to 30 kDa that were detected in both cellular and supernatant fractions of the wild-type and Δ*sphB1* strains (Fig. 1D, lower right panel). These fragments of variable length corresponded to C-terminal degradation products of the FhaB ‘prodomain’ encompassing the proline-rich region (PRR) and CT. The fully processed ∼12-kDa CT was detected exclusively in the wild-type strain culture supernatant, indicating that final processing of the PRR-CT fragment occurs extracellularly and requires the SphB1 protease.

### CT does not affect FhaB folding and surface presentation

It has been previously proposed that CT deletion results in misfolding and loss of biological function of the MCD segment of FhaB (28). We therefore first used negative-stain electron microscopy (EM) to compare the overall structure of FHA molecules purified form culture supernatants of the wild-type (*fhaB*^+^) and *fhaB*ΔCT strains. As shown in Fig. 2A, processing of native FhaB and FhaBΔCT precursors yielded indistinguishable rod-like particles of approximately 4 nm in diameter and ∼45 nm in length, each capped by a globular head at one end. We next employed cryogenic EM (cryo-EM) to compare the filamentous structures formed by FhaB and FhaBΔCT proteins on the surface of *B. pertussis* cells. For this purpose, we analyzed minicells generated by an *ftsQ*-*ftsA*-*ftsZ*-overexpressing strain of *B. pertussis* that also lacked the SphB1 protease and was devoid of fimbriae due to deletion of the *fimABCD* gene cluster (Δ*sphB1*/Δ*fimABCD*). This strategy enabled visualization of individual, unprocessed FhaB molecules in the absence of the potentially confounding fimbrial filaments (Fig. S3 and S4). Indeed, while no filamentous structures were detected on minicells derived from the Δ*fhaB*/Δ*sphB1*/Δ*fimABCD* mutant (Fig. S5), the cryo-EM imaging revealed well-resolved and morphologically undistinguishable FhaB filaments on the surface of minicells expressing either full-length FhaB or its FhaBΔCT variant (Fig. 2B). These exhibited a maximal length of ∼39 nm and a thickness of <4 nm, consistent with the proposed parallel β-helix shaft architecture of the N-terminal FHA-1 region of FhaB (35). Some heterogeneity was observed in minicell preparations from both strains, with subsets of filaments displaying additional periplasmic density, reduced length, or bent conformations (Fig. 2B, lower panels). Together, these data demonstrate that CT deletion does not affect FhaB folding, assembly, or surface exposure.

**Figure 2.**
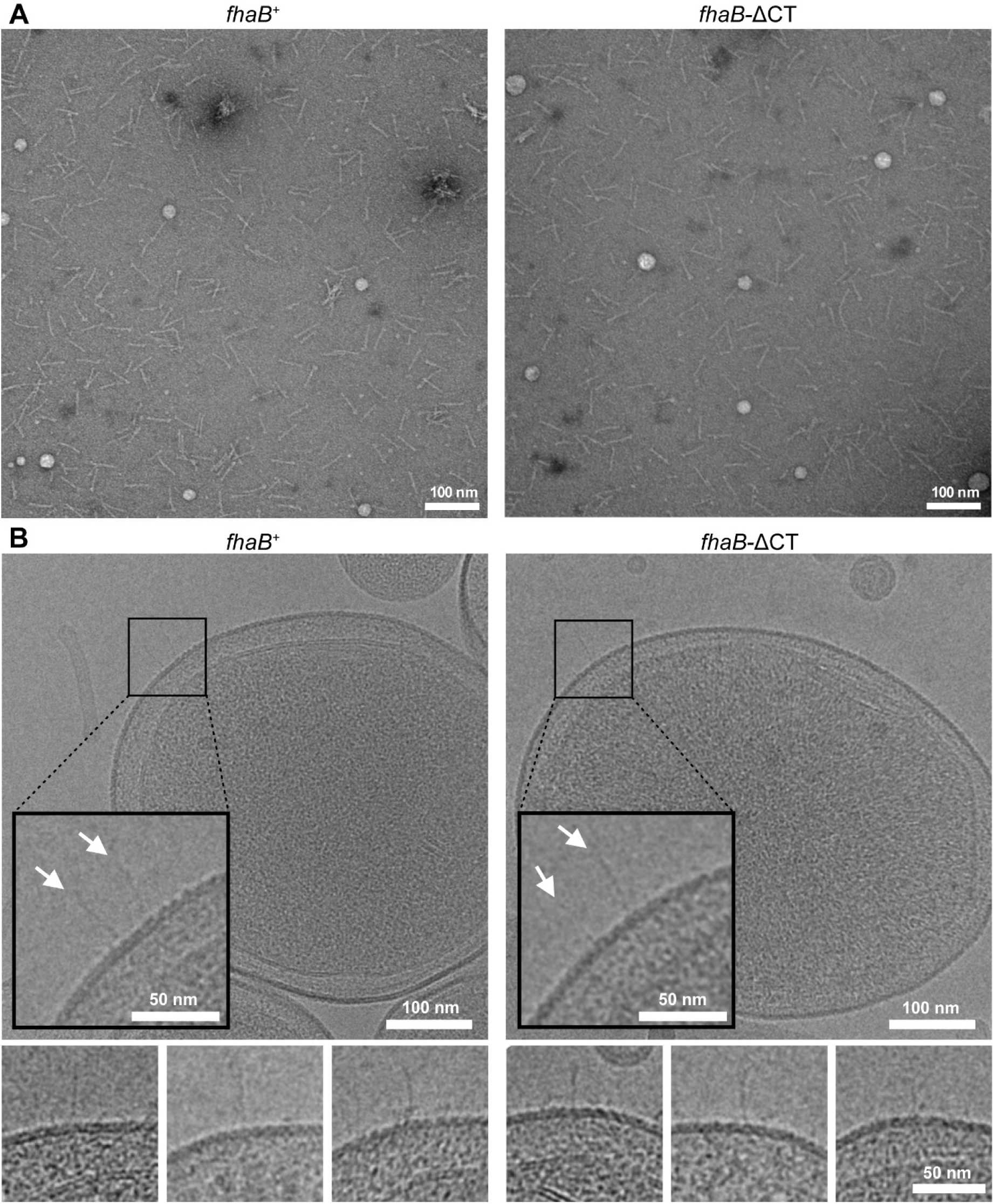
CT deletion does not affect assembly or surface exposure of FhaB. (A) Representative transmission electron micrographs of negatively stained mature FHA particles purified from the culture supernatant of wild-type (*fhaB*, left panel) and *fhaB*-ΔCT (right panel) *B. pertussis* strains. (B) Cryo-electron microscopy images of surface-associated FhaB (*fhaB*, left panel) and FhaB-ΔCT filaments on minicells derived from *B. pertussis* with a Δ*sphB1*/Δ*fimA-D* background. Cell-surface filaments are indicated by white arrows. Lower panels illustrate the pronounced heterogeneity in filament morphology.

### NMR structure of CT

We next determined the solution structure of the CT polypeptide encompassing the C-terminal 95 residues of FhaB (residues 3495–3590) by nuclear magnetic resonance (NMR) spectroscopy. The ^1^H-^15^N heteronuclear single-quantum coherence (HSQC) spectrum exhibited a wide dispersion of backbone amide resonances, consistent with a well-folded protein (Fig. 3A). Conventional three-dimensional triple-resonance and nuclear Overhauser effect spectroscopy (NOESY) experiments provided structural restraints for the majority of residues, enabling assignment of 95% of the backbone and 98% of the side-chain resonances. These restraints were used for structure calculations, and quality statistics for the ensemble of the ten lowest-energy structures (Table 2). As shown in Figure 3B, CT adopts a compact fold comprising an N-terminal α-helix (α_1_) and two twisted β-hairpins (β_1_-β_2_ and β_3_-β_4_) interlinked by a 3_10_ helix (3_10_) and a short β-strand (β_5_). The β-hairpins are positioned on opposite sides of the molecule, generating an apparent pseudo-twofold symmetry when viewed from the top. The α-helix, 3_10_ helix, and β-strands pack tightly via inward-facing hydrophobic side chains, forming a well-defined hydrophobic core. Additional stabilization is provided by hydrogen-bonding interactions between residues located at the termini of the secondary-structure elements. Comparison of the solution structure with the recently solved crystal structure of the CT revealed that the overall protein fold is well preserved but exhibits significant differences in the arrangement of secondary structure elements (Fig. S6). In particular, the β3-β4 hairpin in the crystal structure is markedly extended and twisted toward the N terminus of the CT polypeptide. Moreover, the N-terminal region appears substantially more disordered in the crystal structure, whereas in solution it adopts a well-defined α-helical conformation.

**Figure 3.**
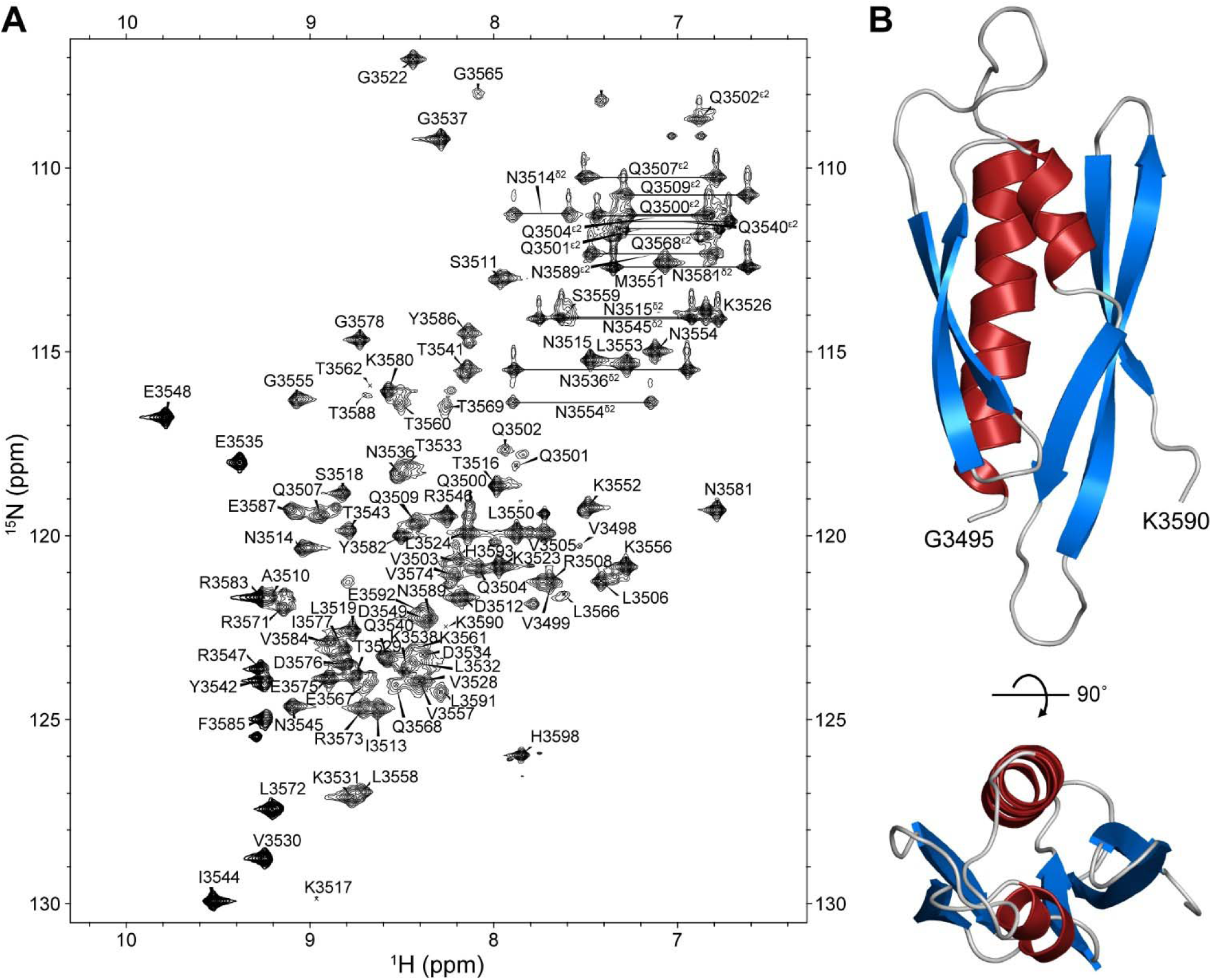
NMR structure of CT. (A) The ^1^H-^15^N heteronuclear single-quantum correlation spectrum of ^15^N-labeled CT (FhaB residues 3495-3590). (B) Cartoon representation of the lowest energy structure of CT, with α-helices shown in red and β strands in blue.

**Table 2.** NMR and refinement statistics for protein structures.

|  |  |
| --- | --- |
| **NMR distance and dihedral constraints** |  |
| Distance constraints |  |
| Total NOE | 2119 |
| Intra-residue | 969 |
| Inter-residue |  |
| sequential ($\vert i - j\vert = 1$) | 347 |
| medium-range ($\vert i - j\vert < 5$) | 259 |
| long-range ($\vert i - j\vert \geq 5$) | 544 |
| Torsion angles restraints |  |
| $\varphi$ | 81 |
| $\psi$ | 81 |
| $\chi$ | 35 |
| **Structure statistics** |  |
| Violations |  |
| distance violations $>0.5 \text{ \AA}$ | 0 |
| torsion angle violations $>5^\circ$ | 0 |
| Average RMS deviation from experimental restrains |  |
| distance restraints ($\text{\AA}$) | $0.010 \pm 0.001$ |
| torsion angle restraints ($^\circ$) | $0.322 \pm 0.056$ |
| Pairwise Cartesian RMSD ($\text{\AA}$) | |
| All/backbone heavy atoms | $3.22 \pm 0.58/2.41 \pm 0.54$ |
| All/backbone heavy atoms without His-tag residues | $1.84 \pm 0.23/1.30 \pm 0.28$ |
| PROCHECK Ramachandran assessment (%) |  |
| Most favored region | $92.30 \pm 1.94$ |
| Additionally allowed region | $7.48 \pm 2.06$ |
| Generously allowed region | $0.22 \pm 0.46$ |
| Disallowed region | 0 |

Although the respective crystal and solution (NMR-derived) structures of the C-terminal domains of *B. pertussis* and *B. bronchiseptica* FhaB are not entirely identical, structural similarity searches revealed that neither of the structures resembles closely any protein deposited in the Protein Data Bank (www.rcsb.org). DALI analysis (56) identified weak similarities to type IV pilus proteins from *Francisella tularensis* (PilE, PDB ID:3SOJ), *Pseudomonas aeruginosa* (PilA, PDB ID:6BBK), and *Neisseria meningitidis* (PilV, PDB ID:5V0M), as well as to the type II secretion system protein GspH from *Escherichia coli* (PDB ID: 2KNQ). However, these matches exhibited low Z-scores (<3.2) and high RMSD values (>3.5 Å). Consistently, PDBeFold analysis revealed only marginal similarity to the type IV-like pilin TTHA1218 from *Thermus thermophilus*, with a Q-score of 0.15 (where 1.0 indicates identical structures). Thus, although CT shares common secondary structure elements with pilin-like proteins, it does not adopt a pilin or pilin-like fold. Instead, CT represents a distinct protein architecture with unique structural features.

### CT plays a crucial role in nasal cavity colonization by *B. pertussis*

We next assessed whether CT contributes to the essential role played by FhaB in the establishment of *B. pertussis* infection of upper airways *in vivo*. C57BL/6J mice were intranasally inoculated with 2×10 CFU in 5-µl doses of suspensions of the indicated bacterial strains and bacterial counts in the nasal cavity were monitored over time. As shown in Fig. 4A, infection with the wild-type (*fhaB*^+^) strain resulted in an approximately two orders of magnitude increase in nasal CFU counts, peaking at day 7 post-infection, followed by a gradual decline at days 14 and 21. In contrast, the *B. pertussis fhaB*ΔCT mutant failed to proliferate on the mouse nasal mucosa, with nasal CFU counts steadily decreasing over time to a near-complete clearance by day 21. Notably, the colonization phenotype of the *fhaB*ΔCT mutant closely resembled that of the Δ*fhaB* strain, indicating that the CT region is essential for sustained *B. pertussis* colonization of the mouse nasal cavity. Importantly, restoration of the intact *fhaB* gene by allelic reintroduction of the deleted CT-coding sequence into chromosome of the *fhaB*ΔCT mutant (*fhaB*ΔCT/CT_in_) restored its nasal colonization capacity to wild-type levels. Hence, the observed loss of virulence of the *fhaB*ΔCT mutant was specifically related to loss of CT function.

**Figure 4.**
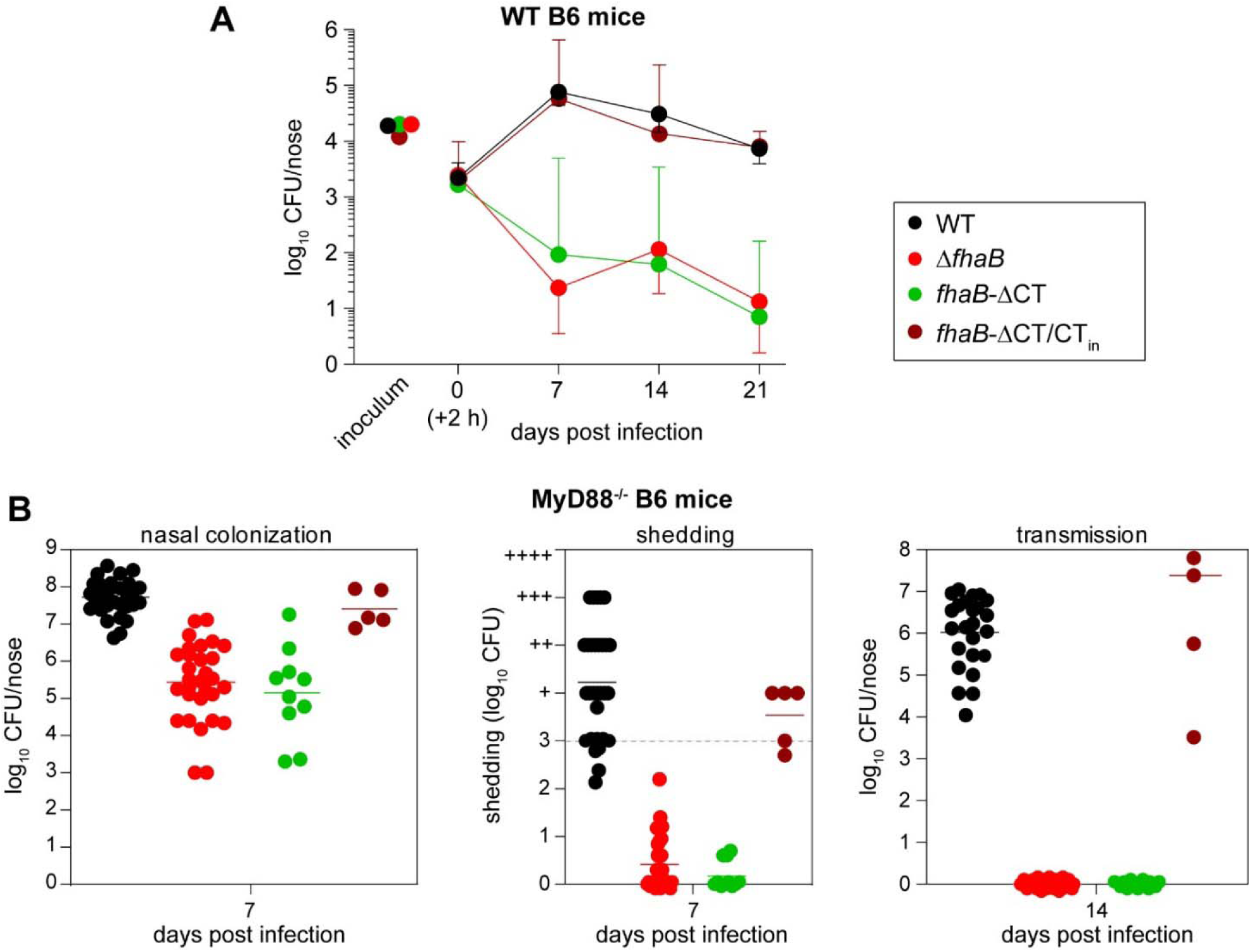
CT plays a crucial role in *B. pertussis* infection and transmission. (A) *B. pertussis* colonization of the nasal cavity in the wild-type C57BL/6J mice following intranasal inoculation (10^4^ CFU in 5 µl) with the indicated *B. pertussis* strains. Each point represents the geometric mean ± standard deviation of bacterial loads (CFU) at the indicated time points, determined by plating homogenized nasal cavity tissues on BG agar and enumerating CFUs after 7 days of incubation at 37 °C from at least 6 animals. (B) Nasal colonization (left panel), bacterial shedding (middle panel) and transmission (right panel) in immunocompromised C57BL/6J MyD88⁻/⁻ mice following intranasal inoculation (10^7^ CFU in 5 µl) with the indicated *B. pertussis* strains. Nasal shedding was assessed after day 7 post infection by gentle tapping of the mouse noses on the surface of a BG agar plate and spreading of deposited bacteria in 100 μl PBS. Shedding above 10³ CFU per mouse nose/plate (dotted line) precluded accurate CFU counting and was therefore estimated from the density of confluent *B. pertussis* lawns on BG agar, scored by (+) signs indicating the approximate order of magnitude. Bacterial transmission was assessed on day 14 by enumerating bacterial loads in the nasal cavities of initially uninfected (recipient) mice following co-housing with infected (donor) mice at a 1:1 ratio for 7 days. Donor mice were removed after 7 days of co-housing. Each point represents the CFU number recovered from a single animal. Horizontal bars indicate geometric means.

To further examine the role of CT in *B. pertussis* shedding and transmission, we employed our recently developed immunodeficient C57BL/6J MyD88⁻/⁻ murine model of human-like catarrhal pertussis infection of the upper airways. The MyD88⁻/⁻ mice lack most of the Toll-like receptor (TLR) signaling, which enables reaching of human-like levels of mouse nasal mucosa colonization by the strictly human-adapted *B. pertussis* bacteria and eventually yields catarrhal pertussis rhinitis (57). Indeed, at 7 days after intranasal inoculation of 10^7^ CFU, the wild-type *B. pertussis* strain reached a colonization density of approximately 10 CFU per nasal cavity, leading to robust catarrhal shedding and efficient transmission of infection to co-housed MyD88⁻/⁻ recipient mice (Fig. 4B). In contrast, infection with the *fhaB*Δ*CT* strain resulted in bacterial burdens approximately three orders of magnitude lower, and comparable to those observed for the *fhaB*-deficient (Δ*fhaB*) strain (Fig. 4B). The reduced colonization was associated with minimal shedding and an absence of transmission to co-housed recipient mice. Collectively, these data demonstrate that CT is critical for FhaB-mediated upper airway colonization by *B. pertussis* and is essential for effective bacterial shedding and transmission to new hosts.

### CT is essential for interaction of *B. pertussis* with ciliated epithelial cells

*B. pertussis* is highly adapted to growth on human upper airway mucosa, where it selectively adheres to ciliated epithelial cells. Such cells can also be obtained by *in vitro* differentiation in an ‘air-liquid interface’ (ALI) culture system (58, 59). To generate such ciliated epithelial layers, we isolated primary human nasal epithelial cells from nasal brushings of anonymous healthy donors and differentiated them on transwell membranes under ALI conditions for at least 28 days. This yielded a continuous, single layer epithelial lining containing ciliated cells (Fig. 5A). Live-cell imaging confirmed active ciliary beating, indicating a well-differentiated and functional airway epithelium that closely mimicked nasal epithelial architecture (Movie S1).

**Figure 5.**
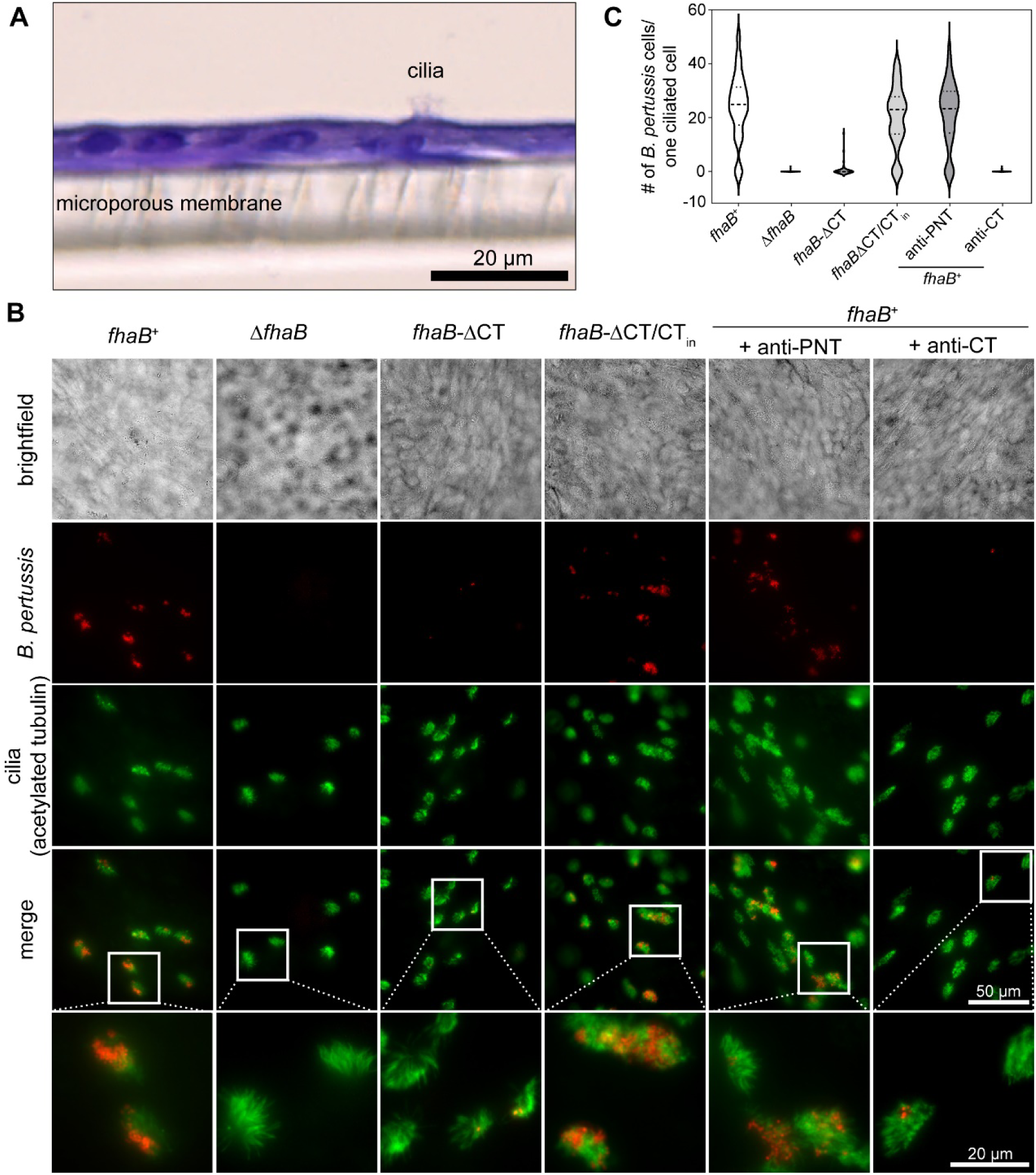
CT is essential for *B. pertussis* interaction with ciliated epithelial cells. (A) Representative image of hematoxylin and eosin-stained paraffin-embedded section of an *in vitro* culture of primary human nasal epithelial cells differentiated for 28 days under ‘air-liquid interface’ (ALI) conditions. Cilia are present on the apical surface of the airway epithelium, whereas the supportive microporous membrane is located on the basolateral side. (B) Fluorescence immunohistochemistry images of ALI-differentiated airway epithelia infected with the indicated *B. pertussis* strains. Cells were infected apically with mScarlet-expressing *B. pertussis* (5×10^5^ CFU in 50 μl of HBSS per transwell) and incubated at 37 °C for 16 h. Following fixation with 4 % (w/v) paraformaldehyde, cells were permeabilized with 0.5 % (v/v) Triton-X100, and cilia were stained using an anti-acetylated tubulin antibody. (C) Violin plot of the numbers of bacteria per one ciliated cell as derived from 50 randomly selected ciliated cells.

The apical surface of ALI-differentiated epithelial layers was inoculated with fluorescent mScarlet-expressing *B. pertussis* strains (5×10^5^ CFU in 50 μl of HBSS). Cell layers were co-cultured with bacteria for 18 hours, washed, fixed, permeabilized and stained for acetylated tubulin prior to assessment of bacterial adhesion by fluorescence microscopy. As shown in Fig. 5B, the wild-type (*fhaB*^+^) strain associated exclusively with ciliated cells and localized within the apical ciliary layer, co-localizing with the acetylated tubulin-stained cilia. Quantitative image analysis revealed that the majority of ciliated cells were colonized, with an average of approximately 25 bacteria associated per ciliated cell (Fig. 5C). In contrast, the Δ*fhaB* mutant failed to adhere to ciliated cells, and no detectable bacteria binding to other epithelial cell types was observed across the transwell surface. Adhesion of the *fhaB*ΔCT mutant was also severely impaired, with only sporadic bacteria detected on a small subset of ciliated cells (Fig. 5B). Importantly, restoration of the CT-coding sequence of the *fhaB* gene on the chromosome (*fhaB*ΔCT/CT_in_) fully restored bacterial adhesion to levels indistinguishable from those of the wild-type strain. These data demonstrate that an intact FhaB, and specifically its CT, is essential for *B. pertussis* adhesion to *in vitro*-differentiated primary human nasal ciliated epithelial cells.

To corroborate the mechanism of CT-mediated adhesion, bacterial binding to ALI-cultured ciliated cells was examined in the presence of anti-PNT or anti-CT sera. As shown in Fig. 5B, anti-PNT serum had no detectable effect on bacterial adhesion, and the number of bacteria associated per ciliated cell was comparable to that observed in the presence of pre-immune serum or in untreated controls (Fig. 5C). In contrast, anti-CT serum effectively blocked bacterial binding to ciliated cells, indicating that CT directly mediates the interaction of *B. pertussis* with ciliated human nasal epithelial cells.

### The C-terminal FhaB moiety remains intact during *B. pertussis* growth on ciliated epithelial cells

The current model of FhaB biogenesis and its processing to FHA is largely derived from *in vitro* studies using bacteria grown in liquid cultures. *In vivo*, however, *Bordetella* cells are not free-floating (planktonic) and instead attach specifically to ciliated epithelial cells, where they proliferate within the ciliary forest in a biofilm-like, host cell-associated environment (60). To examine the fate of FhaB under such physiologically relevant conditions, lysates of *B. pertussis*-infected ALI-differentiated human nasal epithelial cells were analyzed by Western blotting using a panel of antibodies recognizing distinct FhaB segments. Probing with anti-FHA serum revealed limited processing of FhaB to the ∼230-kDa FHA fragment in cilia-adhering wild type bacteria, or to the alternative ∼250-kDa FHA processing product in adhering Δ*sphB1* bacteria (Fig. 6). These FhaB fragments were further N-terminally processed into ∼170-kDa and ∼120-kDa fragments that were detected specifically by anti-MCD serum recognizing the C-terminal segment of mature FHA (Fig. S7).

**Figure 6.**
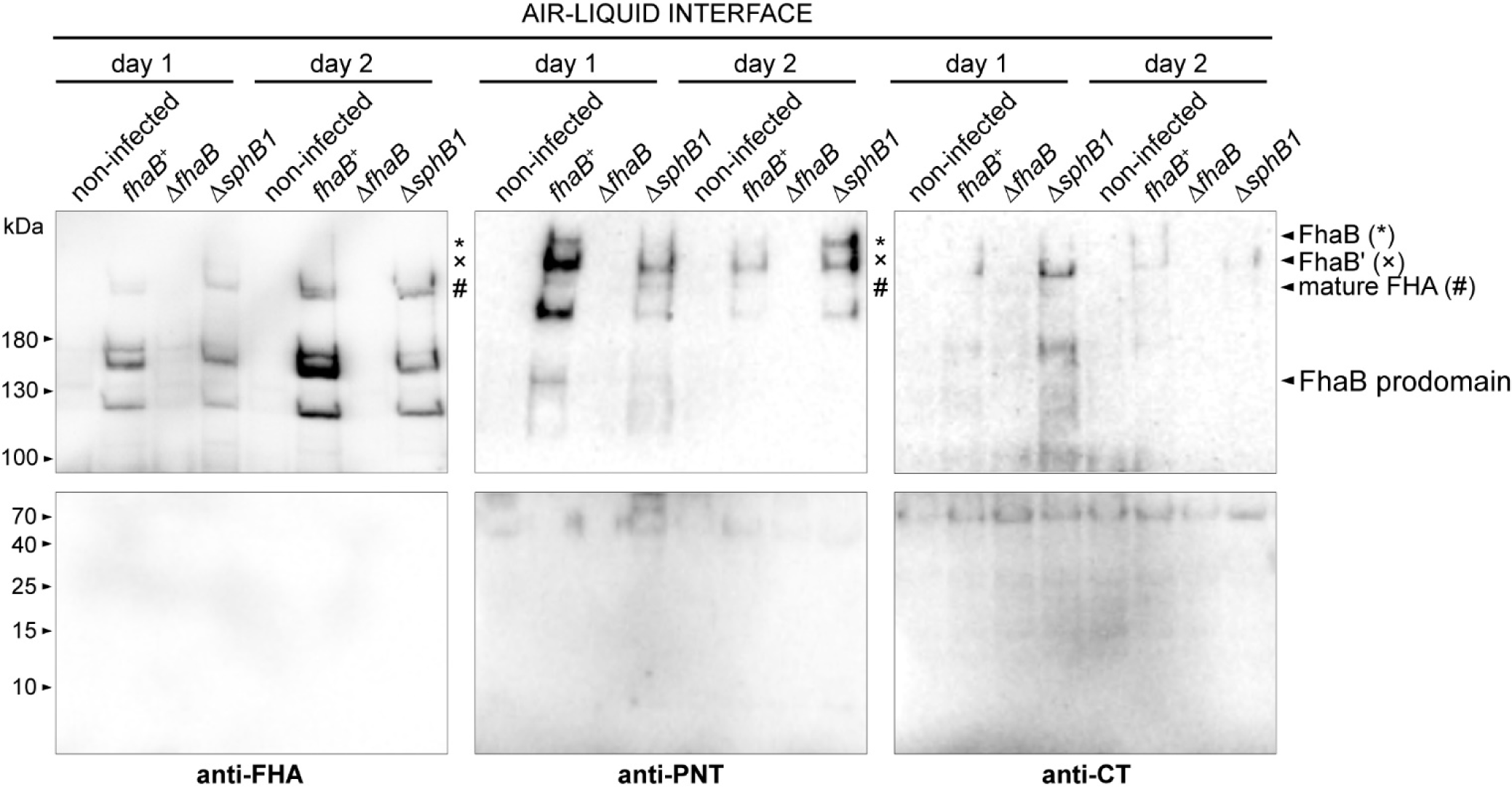
The C-terminal FhaB ‘prodomain’ moiety remains intact during interaction of *B. pertussis* with ciliated epithelial cells. Immunoblot analysis of cell lysates from ALI-differentiated airway epithelial cells that were infected with wild-type or Δ*sphB1 B. pertussis* strains (5×10^5^ CFU in 50 µl HBSS per transwell) for 16 hours and then maintained for an additional 1 and 2 days under ALI conditions. The lysates were separated on 5 % (upper row) and 15 % (lower row) SDS-PAGE gels and membranes containing transferred proteins were probed with anti-FHA, anti-PNT, and anti-CT polyclonal sera. Bands corresponding to distinct FhaB forms are indicated.

In contrast, probing with anti-PNT and anti-CT sera revealed that a substantial proportion of FhaB molecules produced by adhering wild-type (*fhaB*^+^) or SphB1-deficient (Δ*sphB1*) bacteria was not processed to the FHA fragment. Instead, full-length ∼370 kDa FhaB or its N-terminally processed ∼310 kDa variants were prevailing (Fig. 6). Importantly, these FhaB species retained an intact C-terminal ‘prodomain’ moiety, as no lower-molecular-weight (<130 kDa) degradation products were detected by either anti-PNT or anti-CT sera. Together, these data indicate that during adhesion to ciliated epithelial cells, a large proportion of FhaB remains unprocessed and retains an intact C-terminal ‘prodomain’ segment.

### Ultrastructural basis of *Bordetella* attachment to motile cilia

To gain insight into the ultrastructural basis of *Bordetella* attachment to motile cilia at molecular resolution, *in situ* cryo-electron tomography (cryo-ET) was performed on lamellae prepared by cryo-focused ion beam milling of frozen-hydrated samples of infected ciliated cells. In these experiments, the human-pathogenic *Bordetella pertussis* was substituted with the animal pathogen *Bordetella bronchiseptica* that also specifically interacts with ciliated cells and expresses a highly homologous FhaB adhesin. Reconstructed tomograms revealed numerous bacterial cells closely associated with cilia (Fig. 7A). The intact cellular membranes, together with well-preserved axonemal microtubules within the cilia, indicated the high quality of sample preservation. Notably, the tomograms showed that the bacteria do not establish direct membrane-to-membrane contact with cilia; instead, they are typically separated from the cilia by a gap of approximately 50 nm. Closer inspection of the bacterium-cilium interface revealed that the physical contact between the bacterial outer membrane and the ciliary membrane is mediated by filamentous structures that extend from the bacterial surface towards the ciliary membrane (Fig. 7B). The overall morphology of these structures is consistent with FhaB filaments previously observed on the surface of minicells (c.f. Fig. 2B and (55)), suggesting that the physical interaction between bacterial cells and the ciliary membrane is mediated by FhaB molecules. Remarkably, notable differences in filament architecture were observed between the cilia-bound FhaB filaments (i.e., filaments bridging the bacterial surface and the ciliary membrane) and those extending freely (unbound) from the bacterial surface (Fig. 7C). Specifically, cilia-bound filaments were more elongated, with an average length of 55 ± 4.9 nm (Fig. 7D), and exhibited a pronounced increase in electron density at their distal ends, precisely at the point of contact with the ciliary membrane (Fig. 7C). In contrast, unbound filaments were shorter, with an average length of approximately 46 ± 5.3 nm, and were typically characterized by a small globular density at their distal ends (Fig. 7C and 7D). Thus, the differences in FhaB architecture between unbound and cilia-bound filaments likely reflect a structural reorganization of FhaB that is associated with the translocation of the CT from bacterial periplasm upon interaction of the FhaB filament with the ciliary membrane and its subsequent delivery across the ciliary membrane to engage axonemal microtubules.

**Figure 7.**
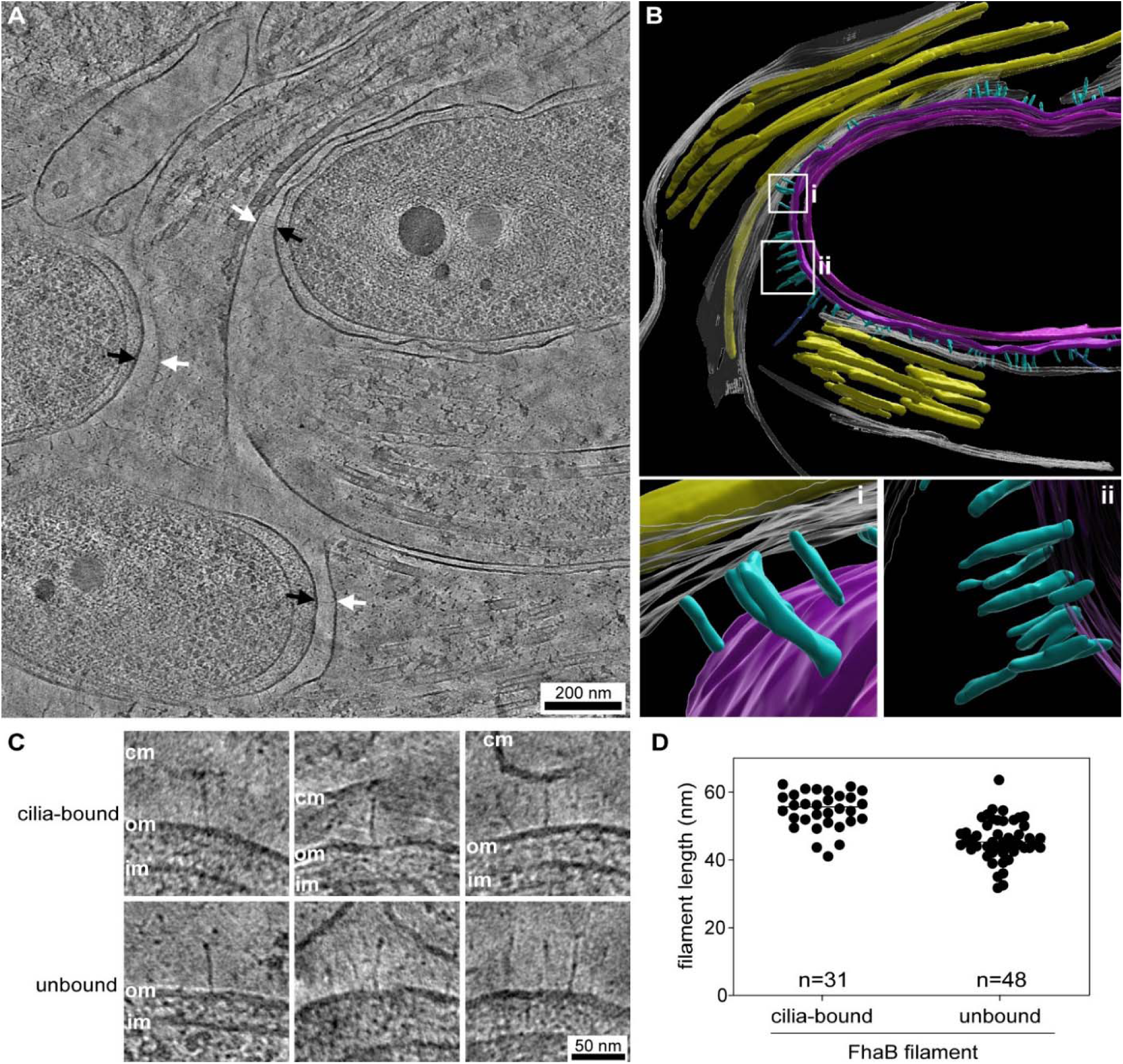
Ultrastructural basis of *Bordetella* attachment to motile cilia. (A) Slice through a cryo-electron tomogram of frozen-hydrated ciliated cells infected with *B. bronchiseptica*. Black arrows indicate the bacterial outer membrane, and white arrows indicate the ciliary membrane. The arrows also highlight FhaB filaments bridging the two membranes. (B) Volume segmentation of the upper-right region of the tomogram shown in (A). FhaB filaments are depicted in turquoise, bacterial envelope membranes in magenta, the ciliary cytoplasmic membranes in white, and axonemal microtubules in yellow. Panels (**i**) and (**ii**) show close-up views of cilia-bound and unbound FhaB filaments, respectively. (C) Close up views of cilia-bound (upper panel) and unbound (lower panel) FhaB filaments. The ciliary membrane (cm), as well as the bacterial outer membrane (om) and inner membrane (im), are indicated. (D) Quantification of the lengths of cilia-bound and unbound FhaB filaments. The number of filaments analyzed in each group is indicated.

## Discussion

We demonstrate that FhaB remains largely unprocessed during *B. pertussis* adhesion to ciliated airway epithelial cells and that the C-terminal domain (CT) of unprocessed FhaB plays a crucial role in the attachment of bacteria to the beating cilia of nasal epithelial cells, thus promoting bacterial colonization of the upper airway mucosa. These findings indicate that full-length FhaB, rather than its so-called ‘mature’ FHA fragment, represents the physiologically relevant adhesin involved in bacterial infection. Accordingly, the extensive production and shedding of the FHA fragment of FhaB observed under liquid *in vitro* culture conditions appears to be a laboratory artifact rather than a biologically relevant process occurring *in vivo*. These results resolve a long-standing paradox in which the ‘mature’ FHA was considered as the active adhesin despite being poorly anchored to the bacterial surface and largely shed into the extracellular milieu, making it difficult to conceptualize how it could sustain host-cell attachment of the bacteria. These results further show that the use of the term ‘prodomain’ for denomination of the ∼1120 residue long C-terminal moiety playing a crucial role in the adhesin function of FhaB is very inappropriate and should be avoided. Indeed, a prodomain is a polypeptide segment that needs to be removed by proteolytic processing that activates a pro-protein, whereas the opposite is the case for FhaB. Its adhesin function depends on the presence of the C-terminal ‘prodomain’ moiety in the unprocessed FhaB that is inactivated by processing to FHA.

The current model of FhaB biogenesis was derived largely from studies with bacteria grown in liquid media and proposes that the C-terminal ‘prodomain’ moiety of FhaB is retained in the periplasm and is rapidly degraded upon SphB1-mediated processing (26)(*c.f.* Fig. 1B). In contrast, our proteomic and immunochemical analysis shows that the C-terminal FhaB moiety is not completely degraded even under *in vitro* liquid culture conditions. Wild-type *B. pertussis* accumulated an ∼130-kDa fragment comprising the entire so-called ‘prodomain’, and this fragment was also recovered in culture supernatants (*c.f.* Fig. 1D). Given that SphB1 is a surface-exposed, outer-membrane-anchored protease that only cleaves extracellular substrates (32), these findings suggest that SphB1 may act on FhaB polypeptides that were translocated into the extracellular milieu. This C-terminal fragment was subject to further proteolytic degradation to produce a stable ∼25-kDa fragment that was unambiguously identified as the C-terminal region of the ‘prodomain’ encompassing the PRR and CT (PRR–CT) (Fig. 1D). Moreover, we detected a standalone CT fragment in culture supernatants of the wild-type (*sphB1*^+^) bacteria, suggesting that the protease-resistant CT domain represents the terminal product of the C-terminal moiety degradation pathway in *B. pertussis* grown in liquid cultures. Such stability of CT is supported by the structural analysis demonstrating that it adopts a compact, well-defined fold with a tightly packed hydrophobic core (Fig. 3B). Collectively, these findings argue against the previously proposed classification of CT as a signal targeting the C-terminal ‘prodomain’ for degradation (18, 26). Instead, its compact fold and apparent surface accessibility supports the role of CT as a ligand-binding module, in line with the recent findings by Costello and colleagues (55).

Overall, we found that FhaB processing in *B. pertussis* adhering to ciliated airway epithelial cells differs markedly from that observed in planktonic bacteria grown in liquid media, where FhaB is predominantly processed by SphB1 to generate the shed FHA fragment. In contrast, a substantial proportion of FhaB in cilia-adhering *B. pertussis* bacteria remains unprocessed, irrespective of the presence of SphB1 (*cf.* Fig. 1 and Fig. 6). These observations suggest that SphB1-mediated FhaB processing is dispensable and does not substantially contribute to the ability of *B. pertussis* to interact with ciliated epithelial cells. Indeed, the SphB1-deficient strain of *B. bronchiseptica* (Δ*sphB1*) showed no detectable defect in colonization of the ciliated epithelium in rat trachea explants (55). Moreover, neither the C-terminal moiety nor its degradation products, such as PRR-CT and CT, were detected in lysates of infected ALI-differentiated epithelial cells, confirming that the C-terminal segment remains intact and escapes extensive proteolysis during epithelial cell adhesion of *B. pertrussis* (Fig. 6). These findings further suggest that contact with ciliated cells stabilizes the full-length FhaB protein and promotes translocation of its C-terminal moiety into the extracellular space, thus protecting it from periplasmic proteases. Alternatively, the cilia-adhering bacteria may represent metabolically adapted cells undergoing a transition from a planktonic to a biofilm-like lifestyle, a shift that could be accompanied by altered protease activities (61). Indeed, biofilm-like lifestyle is thought to represent the dominant mode of *Bordetella* growth during upper airway infection (62).

Of particular importance, in contrast to planktonic bacteria, where processing events are confined to C-terminal regions of FhaB, in the cilia-adhering *B. pertussis* bacteria the processing of FhaB occurs preferentially at its N terminus. This is evidenced by the accumulation of prominent ∼310-kDa and ∼170-kDa degradation fragments, where the former represents a truncated form of full-length FhaB, while the latter corresponds to truncated variants of the FHA fragment, generated by removal of approximately 60 kDa from the N terminus (Fig. 6). Notably, these N-terminal processing events appear to be highly specific and to occur at a single apparent site within the N-terminal segment of the FhaB filament, a region characterized by its pronounced structural stability and resistance to proteolysis under *in vitro* liquid culture conditions (63). As the N-terminal processing activity was not detected in planktonic cultures and was observed exclusively in bacteria associated with ciliated cells, it is likely mediated by a protease of the epithelial cells. Given that the FHA-1 region is critical for the overall stability of the FhaB filament, it is tempting to speculate that such proteolytic processing modulates bacterial attachment dynamics. Specifically, controlled N-terminal cleavage of the FhaB filament may facilitate transient disengagement of bacteria from the ciliary membrane, thus enabling bacterial sliding along the cilia toward the base of the ciliary forest (55). Alternatively, this mechanism may promote the release of individual bacteria from established microcolonies into planktonic state, thereby facilitating dissemination.

A key finding of this study is that the CT of FhaB is essential for *B. pertussis* attachment to nasal ciliated epithelial cells. Deletion of the CT had no discernable effect on FhaB folding, secretion, or surface presentation (Fig. 2), underscoring its critical role in mediating host-pathogen interactions rather than in FhaB biogenesis. This conclusion is further supported by *in vivo* data demonstrating that CT deletion phenocopies the deletion of the entire *fhaB* gene in murine models of nasal *B. pertussis* infection, establishing that the CT is also indispensable for efficient colonization of the upper airway mucosa (Fig. 4). Consistently, a pronounced colonization defect in the murine lower respiratory tract was previously observed for a CT-deficient *B. bronchiseptica* strain (54). This suggests that the requirement for CT function is conserved across *Bordetella* species pathogenic to mammals. Remarkably, we further found that the attachment of *B. pertussis* to ciliated epithelial cells was specifically blocked by anti-CT antibodies (Fig. 5B). Since IgG molecules (∼170 kDa) cannot penetrate the outer bacterial membrane to access the periplasm, these antibodies must recognize an extracellularly exposed CT domain. This finding indicates that the CT is displayed on the bacterial surface rather than remaining confined to the periplasm. This conclusion is reinforced by our observation that FhaB remains largely unprocessed in cilia-adhering *B. pertussis* cells, in line with the translocation of CT to the bacterial surface mediated by the C-terminal FhaB moiety (55). These results make the CT domain to a candidate antigen to be tested for inclusion into future mucosal and parenteral pertussis vaccines, as eliciting of CT-recognizing antibodies could block *B. pertussis* adhesion to airway epithelial cells and thereby improve protection form infection.

The structure and function of FhaB closely resembles that of the antibacterial CdiA proteins of contact-dependent inhibition (CDI) systems (64). CdiA effectors are exported onto bacterial surface by CdiB, which is a β-barrel outer membrane transporter homologous to FhaC (65, 66). During biogenesis, the secretion arrest (SA) domain of CdiA halts CdiB-mediated export such that the C-terminal half of the CdiA protein is retained within the periplasm (67). Thus, FhaB and CdiA share the same unusual surface topology with their N-terminal FHA-1 domains projecting as thin filaments from the bacterium (Fig. 2B and (67)). The distal tip of the CdiA filament is capped by a globular receptor-binding domain (RBD), which recognizes specific cognate receptors on neighboring target bacteria (68–70). CdiA export resumes upon binding its receptor, and the C-terminal effector domain uses the FHA-2 domain to penetrate the target cell envelope and deliver a cytotoxic C-terminal nuclease domain (67). We recently showed that FhaB from *B. bronchiseptica* uses the same overall mechanism to deliver its CT domain into host cells (55). However, rather than delivering a cytotoxic payload, the CT of *B. bronchiseptica* FhaB is a microtubule-binding domain that promotes bacterial colonization through interactions with the ciliary axoneme. These findings are fully consistent with our observations with *B. pertussis* FhaB. Given the high degree of sequence identity between FhaB proteins from *B. pertussis* and *B. bronchiseptica* and the complete conservation of the CT domain sequence across the *Bordetella* genus, we propose that this mechanism is broadly conserved. Thus, the *Bordetella* FhaB proteins appear to represent an evolutionary adaptation of an interbacterial competition system that was repurposed to mediate tight adhesion to host cells (55).

In summary, our findings revise the current model of FhaB biogenesis and provide a mechanistic explanation for the essential role of the CT domain of the FhaB adhesin in *Bordetella* infection. By analogy with the CDI systems, we propose that FhaB is translocated through FhaC and co-secretionally folded into an extended filament displayed on the bacterial surface (Fig. 8). The MCD then acts as the receptor-binding domain (RBD) that recognizes an as-yet-unidentified ligand on the ciliary membrane, thus triggering export of the FHA-2 domain. This would serve as a translocon in the ciliary membrane and enable delivery of the CT domain into the cilium to engage the axonemal microtubules. The FHA-2 translocon is predicted by AlphaFold to form a wing sail-like structure (Fig. 8) and the structural features of this model indeed closely correspond to the pronounced electron density observed as an extension of FhaB filaments at their point of contact with the ciliary membrane (Fig. 7C, upper panel). We thus hypothesize that unlike unbound FhaB filaments, in which the C-terminal moiety of FhaB remains within the periplasm, the cilia-bound FhaB filaments represent the full-length and exposed, ciliary membrane–integrated, form of the FhaB polypeptide that mediates the bacterial attachment to the ciliary membrane (Fig. 8). Collectively, these data redefine FhaB as a multireceptor adhesin and virulence factor whose biological activity extends well beyond serving as the precursor of a shed FHA fragment.

**Figure 8.**
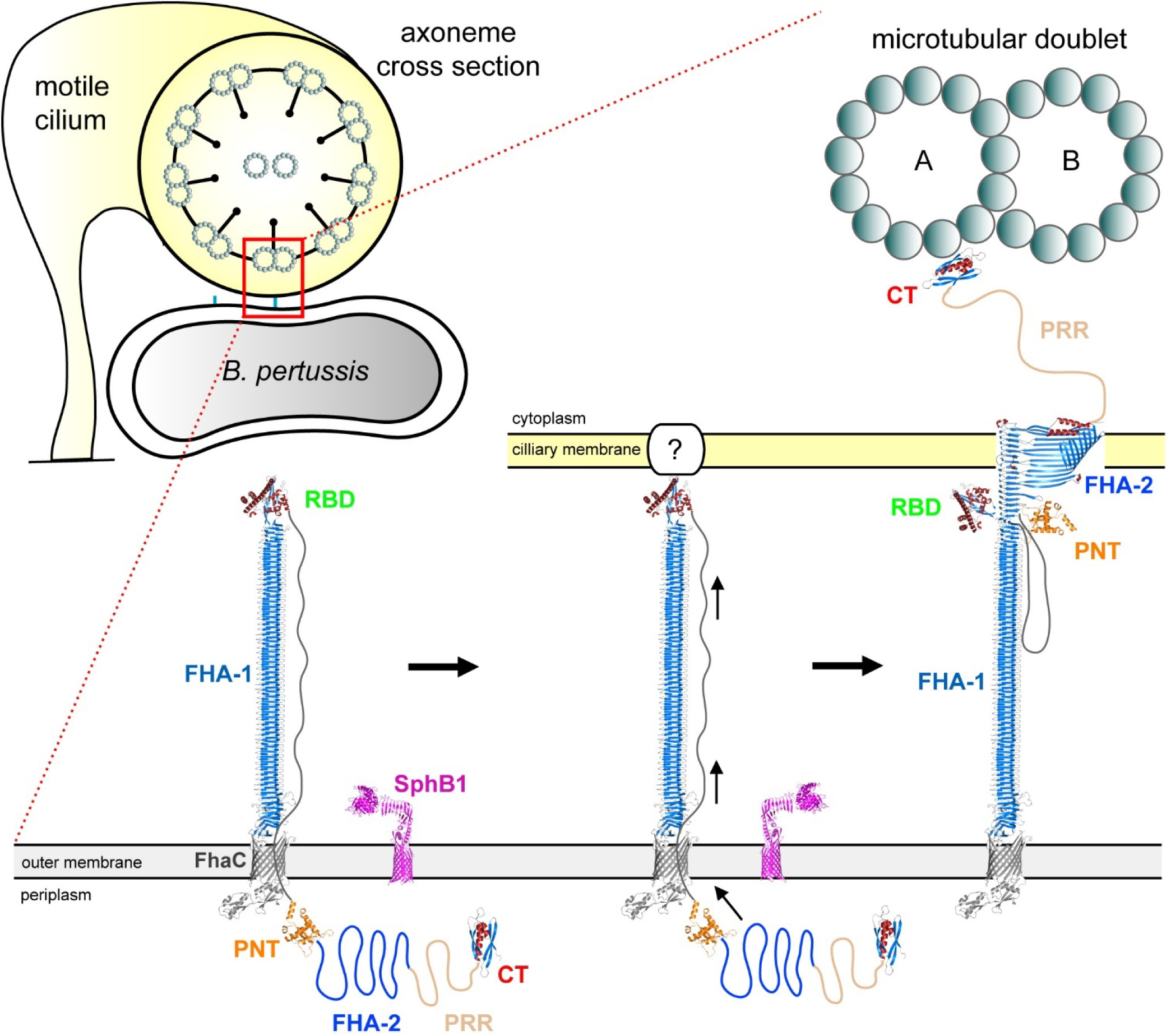
Proposed model of FhaB-mediated *Bordetella* attachment to ciliated epithelial cells. Translocation of FhaB across the outer membrane is mediated by the cognate TPS pore FhaC, from which FhaB emerges in an N- to C-terminal hairpin configuration. Tandem repeats within the FHA-1 domain then sequentially fold into a β-helical shaft, generating an elongated filament capped by a globular receptor-binding domain (RBD aka MCD). Upon RBD interaction with an as-yet-unknown receptor on the cilia of ciliated epithelial cells, FhaB secretion continues, enabling folding of tandem repeats within the FHA-2 domain that extends the FHA-1 domain. The FHA-2 domain is predicted by AlphaFold to form a β-helical shaft decorated with extended β-strand loops, which are proposed to insert into the ciliary membrane and form a conduit for translocation of the C-terminal domain of FhaB (CT). The CT, linked via an unstructured prolyl-rich region (PRR), subsequently engages axonemal microtubules, thereby mediating stable and sustained interactions between *B. pertussis* and host ciliated epithelial cells.

## Materials and Methods

### Ethics statement

All animal experiments were approved by the Animal Welfare Committee of the Institute of Molecular Genetics of the Czech Academy of Sciences, Prague, Czech Republic. Handling of animals was performed according to the Guidelines for the Care and Use of Laboratory Animals, the Act of the Czech National Assembly, Collection of Laws no. 246/1992. Permissions no. 1989/2023 was issued by the Animal Welfare Committee of the Institute of Molecular Genetics, Czech Academy of Sciences, Prague, Czech Republic. Sampling of nasal swabs was performed from healthy anonymous donors using a non-invasive procedure and was approved by the Ethics Board of Thomayer Hospital in Prague, Czech Republic, on April 24, 2024 (Docket No. 10358/24; G-24-10).

### Antibodies and reagents

The suicide vector for allelic exchange on the *B. pertussis* chromosome (pSS4245) was provided by Scott Stibitz (FDA, Center for Biologics Evaluation and Research, Silver Spring, USA). The rabbit polyclonal anti-FHA antibody was gift of Branislav Vecerek (Institute of Microbiology, Prague, Czech Republic).

### Bacterial strains and growth conditions

The *Escherichia coli* XL1-Blue was used for plasmid construction and the *E. coli* SM10 λpir was used for bacterial conjugation. *E. coli* strains were cultivated at 37°C on Luria-Bertani (LB) agar in the presence of ampicillin (150 μg/ml). *Bordetella pertussis* Tohama I (CIP 81.32) and *Bordetella bronchiseptica* RB50 were cultivated on Bordet-Gengou (BG) agar plates (Difco, USA) supplemented with 15 % defibrinated sheep blood (LabMediaServis, Czech Republic). For mice experiments, a spontaneous streptomycin-resistant *B. pertussis* strains were isolated from the BG agar plates supplemented with 1% glycerol, 15% defibrinated sheep blood and 100 μg/ml streptomycin. Liquid cultures of *B. pertussis* were grown in a modified Stainer–Scholte (SS) medium supplemented with 5 g/l of casamino acids and 1 g/l of heptakis (2,6-di-O-dimethyl)-β-cyclodextrin. The fluorescent strains of *B. pertussis*, carrying the pBBR1-mScarlet plasmid (71) were cultivated on solid or liquid media in the presence of 10 μg/ml chloramphenicol.

### Plasmid construction

The pET22b-CT vector was prepared by ligation of the NdeI/XhoI-digested pET22b with the NdeI/XhoI-cleaved PCR product amplified from genomic DNA of *B. pertussis* using the forward 5’-TTTCATATGGGGCGCCACGTGGTGCAA-3’ and the reverse 5’-TTTCTCGAGTTTGTTGGTTTCATAGAAGACC-3’ primers. The pET22b-PNT vector was constructed by cloning of the PCR product amplified from genomic DNA of *B. pertussis* using a pair of primers (5’- TGGATATCGGAATTAATTCGACGGTCCGGGTCGCGCC-3’ and 5’-GTGGTGGTGGTGGTGCTCGAGGGCGTTCCCCTTCGCG-3’) into the BamHI/NotI-digested pET22b using the Gibson assembly strategy. The pET22b-MCD vector was prepared by ligation of BamHI/XhoI-digested pET22b plasmis with the PCR fragment amplified using a pair of primers (5’-TGGATATCGGAATTAATTCGGCCGATGCGCGCAAGGAC-3’ and 5’-GTGGTGGTGGTGGTGCTCGAGTTTGCTCTGGTCGATAAACTTG-3’) using the Gibson assembly strategy. For construction of the pSS4245-*fhaB*Δ*CT* allelic exchange vector, two DNA fragments corresponding to the 5′ and the 3′ flanking regions of the in-frame deletion of codons 3493-3590 of the *fhaB* gene were amplified from genomic DNA of *B. pertussis* using two pairs of primers (5’-CCTATGCTAGGGCGGCCGCAAAGCACTGGGCCGGAGGC-3’, 5’-CGGCAGGCCGCGACTACCTAGGGCGTCGTTTTGCCCGGC-3’ and 5’- GCCGGGCAAAACGACGCCCTAGGTAGTCGCGGCCTGCCG-3’, 5’-AGGACGCGTGGATCCGAATTGGAGAACCCCACCTGACTG-3’) and into the SpeI/EcoRI-digested pSS4245 vector using the Gibson assembly strategy. The pSS4245-ΔCT/CT_in_ vector was constructed by cloning of the PCR fragment amplified from genomic DNA of *B. pertussis* using the forward 5’-CCTATGCTAGGGCGGCCGCAAAGCACTGGGCCGGAGGC-3’ and the reverse 5’- AGGACGCGTGGATCCGAATTGGAGAACCCCACCTGACTG-3’ primers into the SpeI/EcoRI-digested pSS4245 vector using the Gibson assembly strategy. The pSS4245-*sphB1* allelic exchange vector was prepared by ligation of the SpeI/NdeI- and NdeI/EcoRI-digested DNA fragments corresponding to the 5′ and the 3′ flanking region of the in-frame deletion of the *sphB1* gene amplified from genomic DNA of B. pertussis by using two pairs of primers (5’-TTTACTAGTAACAACGCTTCATAGCAGCATA-3’, and 5’-TTTCATATGGGGCGGGGGAGCCTCC-3’; and 5’-TTTCATATGCCCTGCGCCCGCCTCG-3’ and 5’-TTTGAATTCTCCATCGGGTCGATCAGCA-3’) into the SpeI/EcoRI-digested pSS4245 vector. The pBBR1-*P*_ara_-*ftsQAZ* was generated by Gibson assembly of PCR fragments containing the *E. coli* arabinose *(ara)* promoter (amplified with 5’-GCTGGGTACCGGGCCCCCCCCAATTGTCTGATTCGTTACC-3’ and ATGGTGCGAGCGTCGTTCCACATATGTATATCTCCTTCTTAA-3’ primers), and *B.pertussis ftsQAZ* operon (amplified with 5’-TGGAACGACGCTCGCACCAT-3’ and 5’-CGGTGGCGGCCGCTCTAGAATCAATCGGCTTGCTTGCGCA-3’ primers) with SpeI/XhoI-cleaved pBBR1 plasmid. All constructs were confirmed by DNA sequence analysis with an ABI Prism 3130XL analyzer (Applied Biosystems, USA) using a BigDye Terminator cycle sequencing kit.

### Mutagenesis of *B. pertussis*

Mutant *B. pertussis* strains were constructed by homologous recombination using the suicide allelic exchange vector pSS4245 as described previously (57). In brief, the *B. pertussis* cells grown on BG agar supplemented with 50 mM MgSO_4_ were mated at 37°C for 6 h on fresh BG agar plates supplemented with 50 mM MgSO_4_ and 50 mM MgCl_2_ with *E. coli* SM10 λpir harboring the respective pSS4245 construct. The transconjugants were selected on BG agar supplemented with 50 mM MgSO_4_, 500 μg/ml streptomycin, 30 μg/ml ampicillin and 40 μg/ml kanamycin. Individual colonies were streaked on BG agar and the resulting single colonies were characterized for the presence of desired mutation by restriction analysis of PCR fragments obtained from colony PCR and sequencing of relevant PCR-amplified segments.

### Protein purification

The MCD, PNT and CT domains of FhaB were expressed as recombinant C-terminally His-tagged proteins in *E. coli* strain BL21(λDE3) transformed with the appropriate plasmid. Exponential 500-ml cultures were grown in a shaking incubator at 37 °C in MDO medium (yeast extract, 20 g/l; glycerol, 20 g/l; KH_2_PO_4_, 1 g/l; K_2_HPO_4_, 3 g/l; NH_4_Cl, 2 g/l; Na_2_SO_4_, 0.5 g/l; and thiamine hydrochloride, 0.01 g/l) supplemented with 150 μg/ml of ampicillin. For NMR experiments, the cells were grown in M9 medium containing 1 g/l ^15^NH_4_Cl (Cambridge Isotope Laboratories, USA) and 2 g/l [^13^C]glucose (Cambridge Isotope Laboratories, USA). Expression of proteins was induced by adding 1 mM isopropyl-β-d-thiogalactopyranoside (IPTG, Alexis, Switzerland) at an optical density at 600 nm (OD_600_) of 0.6. After 4 additional hours of cultivation at 37 °C, the cells were washed with PBS and disrupted by a Misonix S-4000 ultrasonic processor (Misonix, USA) at 4 °C. The cell lysates were centrifuged at 3,000 g for 5 min to remove unbroken cells and the supernatant was centrifuged at 20,000 g for 30 min at 4 °C. For purification of CT, the supernatant was loaded on Ni Sepharose 6 High Performance column (Cytiva, USA) equilibrated with PBS supplemented with 150 mM NaCl (PBSN), washed extensively with PBSN supplemented with 50 mM imidazole, and the CT protein was eluted from the column with PBSN buffer supplemented with 200 mM imidazole. The collected fractions were concentrated using an Amicon YM3 ultrafiltration membrane (Millipore, USA) and the concentrate was loaded on a Superdex 75 gel filtration column (Cytiva, GE Healthcare, USA) equilibrated with PBS. The collected fractions were pooled, concentrated with an Amicon YM3 ultrafiltration membrane, and stored at -20 °C. For purification of MCD and PNT, pelleted inclusion bodies were solubilized with buffer containing 50 mM Tris-HCl (pH 8.0) and 8 M urea (TU) and the urea extract was clarified by centrifugation at 20,000 g for 30 min at 4 °C. The supernatant was loaded on Ni Sepharose 6 High Performance column (Cytiva, USA) equilibrated with TU buffer, the column was washed with TU buffer supplemented with 20 mM imidazole and the PNT protein was eluted with TU buffer containing 100 mM imidazole. FHA fragments were purified as previously described (63). Briefly, *B. pertussis* cultures (600 ml) grown in modified SS medium supplemented with 10 g/l casamino acids for 40 h at 37 °C were centrifuged at 14,000 g for 25 min at 4 °C. The resulting supernatants were filtered through a 0.22-µm TPP Filtermax rapid bottle-top filter (TPP, Switzerland). Clarified supernatants were loaded onto 800 µl of Cellufine Sulfate resin (JNC Corporation, Japan) pre-equilibrated with 10 mM PBS. After washing with PBS, FHA proteins were eluted with 10 mM phosphate buffer supplemented with 700 mM NaCl. The purity of all protein preparations was monitored by SDS-polyacrylamide gel electrophoresis (SDS-PAGE), and protein concentrations were determined by Bradford assay (Bio-Rad) using bovine serum albumin as a standard.

### Preparation of mouse polyclonal sera

Five-week-old female BALB/cByJ mice (Charles River, France) were immunized with intraperitoneal injection of the MCD, CT and PNT proteins (30 μg per mouse) formulated with an incomplete Freund’s adjuvant (Sigma). Control mice were vaccinated with the adjuvanted PBS. Mice received three doses of the vaccines in two-weeks interval and one week after the third immunization blood was collected from anesthetized animals (i.p. injection of 80 mg/kg ketamine and 8 mg/kg xylazine) by retroorbital puncture method. Sera were recovered from the supernatant after centrifugation of clogged blood at 5,000 g for 10 min at 8 °C and stored at -20 °C.

### Mouse infection, bacterial shedding and transmission

Nasal colonization, bacterial shedding and transmission were determined as described previously (57). In brief, for determination of long-term nasal colonization, female C57BL/6J mice (Charles River, France) were anesthetized by intraperitoneal injection of ketamine (80 mg/kg) and xylazine (8 mg/kg) in 0.9 % saline and challenged intranasally with 5 μl of 1×10^4^ CFU of *B. pertussis* cells. At indicated time points, mice were sacrificed and the dissected nasal cavities with turbinates were homogenized in PBS using an IKA Ultra Turrax T25 tissue homogenizer (Sigma-Aldrich, USA). The suspension was cleared of bone debris by centrifugation at 217 g for 10 min and serial dilutions of the supernatant were plated on BG agar supplemented with 15% defibrinated sheep blood and 100 μg/ml of streptomycin. The CFU counting was carried out after incubating the plates at 37 °C for 4 days. Bacterial shedding and transmission experiments were performed with in-house-bred C57BL/6J MyD88^-/-^ mice (MyD88 null, B6.129P2(SJL)-Myd88tm1.1Defr/J, Jackson Laboratory, USA) challenged intranasally with 5 μl of 1×10^7^ CFU of *B. pertussis* cells (57). Shedding of bacteria from the nose of the infected mice was quantified on day 7 by gentle tapping of infected mouse noses four times on the surface of a BG agar plate containing 15% sheep blood and 100 μg/ml of streptomycin. The recovered bacteria were supplemented with 100 μl of PBS and spread over the plates. The CFU counting was carried out after incubating the plates at 37 °C for 4 days. For transmission experiments, intranasally infected index (donor) mice were co-housed in the same cage with non-infected (recipient) mice at a ratio of 1:1. On day 7, the bacterial shedding of donor mice was assessed before the donor mice were sacrificed, and bacterial loads in their nasal cavity were determined as described above. The recipient mice were kept for additional 7 days (14 days after exposure to infected mice) before being sacrificed. The bacterial loads in their nasal cavity were then determined as described above.

### Air-liquid interface (ALI) cultures of human nasal epithelial cells

Cultures of human nasal epithelial cells were prepared as previously described (72). Briefly, nasal epithelial cells were collected from the anterior nares of anonymous healthy donors using a cytology brush. Cells were digested with TrypLE Select (Gibco, USA) for 5 min at 37°C and single-cell suspensions were centrifugated at 350 g for 5 min. The pelleted was gently resuspended in nasal epithelial cell (NEC) medium and seeded into T25 flasks containing mitomycin-treated 3T3-J2 fibroblasts in NEC medium. After expansion, 5×10^4^ cells were plated onto the apical side of collagen-coated 6.5-mm Transwell inserts with 0.4 μm pore polyester membrane (Corning, USA) in 200 μl apical and 600 μl of basolateral NEC medium without fungin. After 72 h, when cells reached confluency, the apical medium was removed to establish air-liquid interface conditions (air-lifting), and the basolateral medium was replaced with differentiation ALI medium. The ALI medium was refreshed three times per week. The day of air-lifting was designated as day 0 of the ALI culture, and the experiments were conducted between days 25 and 28. To prevent mucus accumulation, the apical surface was washed with PBS for 30 min every 3 days starting from day 14. Differentiation and epithelial integrity were assessed by visual inspection and measurement of transepithelial electrical resistance (TEER) prior to each experiment. Additionally, on day 23, the basolateral ALI medium was replaced with antibiotic-free ALI medium, with at least two medium changes performed before experimentation.

### Histological analysis

ALI-differentiated cells were immersed in 4 % (w/v) paraformaldehyde in PBS at 4 °C for 24 h, followed washing with PBS and transfer to 70 % ethanol solution for 24 hours at 4°C. Tissue samples were processed using a Leica ASP6025 Tissue Processor (Leica Biosystems, Germany) and embedded in paraffin blocks using a Leica EG1150 Modular Tisue Embedding Center (Leica Biosystems, Germany). Blocks were sectioned at 2 μm using a Leica RM2255 Fully Automated Rotary Microtome (Leica Biosystems, Germany), and sections were mounted on Superfrost PLUS slides (Thermo Fisher Scientific, USA). Slides were stained with hematoxylin and eosin using a Leica ST5020 Multistainer (Leica Biosystems, Germany) and stained samples were mounted using a Pertex mounting medium (Histolab, Sweden) in a Leica CV5030 Fully Automated Glass Coverslipper (Leica Biosystems, Germany). Images were acquired using an AxioScan.Z1 Slide Scanner (Carl Zeiss, Germany) with 20× objective.

### Bacterial infection of ALI-differentiated cells

Differentiated ALI cultures of human nasal epithelial cells were infected from apical side with mScarlet-expressing *B. pertussis* (5×10^5^ CFU in 50 μl of HBSS) and incubated at 37 °C in a 5 % CO_2_ atmosphere. After 16 h of incubation, excess liquid was removed from the apical side. The cells were then washed and either fixed with 4% (w/v) paraformaldehyde in PBS at 4 °C for 24 h (for fluorescence microscopy) or lysed with 200 µl of lysis buffer containing 50 mM Tris-HCl (pH 7.4), 8 M urea, 1% SDS, and 10 mM DTT (for Western blot analysis). Following fixation, the cells were washed with PBS (3×10 min of incubation) and permeabilized with 0.5 % (v/v) Triton-X100 in PBS at room temperature for 30 min. The permeabilized cells were washed with PBS (3×10 min of incubation) and blocked with 1 % BSA in PBS at room temperature for 1 h. The blocked samples were incubated with a mouse monoclonal anti-acetylated tubulin antibody (Sigma, T6793) at room temperature for 1 h at a dilution of 1:500 in PBS. Following washing step with PBS (3×10 min of incubation), the cells were incubated with AlexaFluor647 F(ab’)2 fragment goat anti-mouse (Jackson ImmnunoResearch, 115-606-072) at room temperature for 30 min at a dilution of 1:500 in PBS. After washing step with PBS (3×10 min of incubation), the dissected membrane was mounted on glass slide (Waldemar Knittel Glasbearbeitungs, Germany) in a Vectashield mounting medium (Vector Laboratories, USA) and sealed with a coverslip (Hirschmann Laborgeräte, Germany).

### Fluorescence microscopy

Fluorescence microscopy was conducted using an Olympus IX83 inverted microscope equipped with a Prime95B camera (Teledyne Photometrics, USA) and cellSens Dimension software (Olympus, USA). Images were magnified using UPLFLN10x2PH (NA 0.3) and UPLXAPO100xOPH (NA 1.45) objectives. The fluorescent signal was acquired using combination of excitation and emission filters: 555/25 and 605/52 for mScarlet, and 645/30 and 705/72 for AlexaFluor647. The images were analyzed using Fiji (v1.54f) software (https://imagej.net). The binding capacity of bacteria was quantified by counting the number of bacteria per single ciliated cell from at least 50 randomly selected ciliated cells. Violin plots were generated using GraphPad Prism v10.0.3. Statistical significance of bacterial count difference between the untreated and the antibody-treated *B*. *pertussis* cells was assessed by one-way ANOVA followed by Dunnett’s multiple comparison test, **** (p < 0.0001).

### Negative-stain electron microscopy

Freshly purified mature FHA proteins were diluted to 1 µg/ml in PBS and 3 µl of the sample was applied onto glow-discharged 200-mesh carbon-coated copper grids (Electron Microscopy Sciences, USA). Excess sample was removed with filter paper and stained with two drops (3 µl) of freshly prepared 2% uranyl acetate (Electron Microscopy Sciences, USA). After blotting the stain, the grids were desiccated at room temperature and imaged using a Jeol JEM-1400 transmission electron microscope equipped with a bottom-mounted FLASH 2k×2k CMOS camera.

### Cryo-electron microscopy

Minicells were generated by overexpression of *ftsZAQ* in the *fhaB, fhaB*-ΔCT and the *fhaB*-deficient strains of *B. pertussis* carrying a Δ*sphB1*/Δ*fimA-D* background. Bacteria were grown in modified SS medium supplemented with 15 μg/ml chloramphenicol at 37 °C to an OD_600_ of 0.2, after which minicell production was induced by the addition of 10 mM L-arabinose for 48 h. Following induction, intact bacteria were removed by centrifugation at 3,000 *g* for 20 min at 4 °C and the minicell-containing supernatant was collected. Minicells were pelleted by centrifugation at 42,000 *g* for 1 h at 4 °C, resuspended in 10 mL of PBS and further purified by centrifugation on a discontinuous 5-20 % OptiPrep gradient (Serumwerk Bernburg, Germany) at 4,600 *g* for 30 min at 4 °C. Isolated minicells were washed twice with PBS by repeated centrifugation steps at 42,000 *g* for 1 h at 4 °C and finally resuspended in 100 µl of PBS. For cryo-electron microscopy, 3 µl aliquots were applied to glow-discharged Quantifoil R 2/1 Cu 200 Mesh holey carbon grids (Quantifoil, SPT Labtech, UK) and plunge-frozen into liquid ethane using a Leica EM GP2 automatic plunge freezer (Leica, Germany). Grids were imaged using either a 200 kV Jeol JEM-F200 transmission electron microscope (Jeol, Japan) equipped with TemCam XF416 cooled CMOS camera (TVIPS, Germany) and Fishione Model 2550 Cryo Transfer Tomography Holder (Fishione Instruments, USA), or a 200 kV Talos Arctica transmission electron microscope equipped with a Falcon 4 direct electron detector (Thermo Fisher, USA).

### Cryo-electron tomography

Differentiated ALI cultures of human nasal epithelial cells were infected apically with 5×10^5^ CFU of the ΔBB0450-0452 mutant strain of *Bordetella bronchiseptica* RB50 carrying the pBBR1 plasmid expressing the mScarlet fluorescent protein (55, 71). Bacteria were applied in 50 μl of HBSS, and the cultures were incubated at 37 °C in a 5 % CO_2_ atmosphere. After 16 h, excess liquid was removed from the apical side of the transwells and the cultures were then incubated for an additional 24 h under the same conditions. Subsequently, the differentiated cell layer was treated with Accutase solution (Sigma, USA) for 1 h at 37 °C in a 5 % CO₂ atmosphere. The resulting single-cell suspensions were washed with HBSS and 3 µl of the suspension was deposited onto glow-discharged holey carbon coated Quantifoil R 2/1 200 mesh copper grids (Quantifoil Micro Tools, Germany). Excess liquid was blotted from the grids using a Leica EM GP2 (Leica Microsystems), and the grids were vitrified by plunging into liquid ethane. Chamber conditions were set to 22 °C and 95 % humidity, with a blotting time of 7 s and a blotting sensor setting of 1 mm extra travel. Preparation of lamelae for cryo- electron tomography was performed according to a modified protocol (73). Briefly, grids were sputter-coated with 20 nm of platinum using a Leica EM ACE600 (Leica Microsystems, Germany) under cryogenic conditions and subsequently inspected using a Leica CryoCLEM Thunder imager (Leica Microsystems, Germany) at a stage temperature of −180 °C. Whole-grid montages were acquired using integrated software with transmitted light and red fluorescence. Regions containing bacteria-decorated ciliated cells near the center of grid squares were selected for Z-stack acquisition. Z-stacks were collected with a step size of 0.33 µm using reflected light and the red fluorescence channel. Fluorescence signals were used to localize bacterial clusters in X, Y, and Z, while reflected light was used to determine the Z-distance between the cluster center and the sample surface. Samples were then transferred to a Ga^2+^ focused ion beam scanning electron microscope (FIB-SEM) Tescan Amber 2 (Tescan Group, Czech Republic) equipped with a Leica cryo stage and a Leica VCT500 cryo transfer system (Leica Microsystems, Germany). Maximum intensity projections generated from fluorescence images, with marked target positions, were overlaid onto FIB-SEM images of grids tilted to 90° relative to the FIB-SEM axis using 3-point affine registration. Fiducial markers were milled adjacent to the regions of interest. The stage was subsequently tilted to 10° relative to the FIB-SEM axis, and the fiducial markers, together with depth information, were used to determine the lamella position. Rough milling (>1 µm from the final lamella) was performed at 1 nA, intermediate milling at 250 pA (∼500 nm from the lamella), and final polishing to a thickness of ∼150 nm was carried out at 25 pA. Grids containing finished lamellae were transferred to a Jeol JEM-F200 transmission electron microscope (Jeol, Japan) equipped with a Gatan Alpine direct electron camera (Gatan, USA) and a Simple Origin Model 210 cryo transfer holder (Simple Origin, USA). Tomographic data were acquired using SerialEM software (74) at a nominal magnification of 10,000×, corresponding to a pixel size of 0.447 nm. Tilt series were collected from −60° to +60° with 3° increments using a dose-symmetric scheme, resulting in a total accumulated dose of ∼90 e⁻/Å² (75). Tomograms were reconstructed using the IMOD software package (76). Reconstructed tomograms were semi-manually segmented using Microscopy Image Browser v2.9102 (77), and the final volumes were rendered in Imaris v11.0.1 (Oxford Instruments, UK).

### Nuclear magnetic resonance (NMR) spectroscopy

All NMR spectra of the CT protein were recorded at 25 °C on the Bruker Avance III 700 MHz spectrometer equipped with the cryogenic ^1^H(^19^F)/^13^C/^15^N TCI probe head. All spectra were acquired using the TopSpin 3.6 and processed with the NMRPipe (78). Spectra were analyzed using NMRFAM–SPARKY 1.470 software (79). The ^1^H chemical shifts were calibrated using the water signal as internal reference and both ^13^C and ^15^N chemical shifts calibrated through indirect referencing to DSS and liquid ammonia, respectively. Backbone assignments were obtained from a set of standard 3D triple-resonance experiments, including HNCACB, CBCA(CO)NH, HNCA, HN(CO)CA, HN(CA)CO, and HNCO. Side-chain assignments were based on the (H)CC(CO)NH, H(CC)(CO)NH and HCCH-TOCSY experiments (used for the assignment of the aliphatic and aromatic side chains). The connectivity between aliphatic and aromatic spin systems was derived using 2D (HB)CB(CGCD)HD and 2D (HB)CB(CGCDHD)HE spectra. The ^15^N-edited and ^13^C-edited NOESY–HSQC (aliphatic and aromatic) spectra with mixing time of 120 ms were recorded to obtain inter-proton distance restraints. The inter-proton distance restrains (Table 1) were obtained from NOE cross peaks from NOESY spectra by ARIA webserver (80). The protein backbone angles φ, ψ, and χ_1_ were derived from the ^1^H^N^, ^1^H^α^, ^15^N, ^13^C′, ^13^C^α^, and ^13^C^β^ chemical shifts using TALOS-N software (81). Structure refinement was performed with CNS version 1.3 using the RECOORD water refinement protocol (82). The quality of the final structures was checked by PROCHECK (83).

### Mass spectrometry

The supernatant (37 ml) obtained after the centrifugation of the late-exponential culture of *B. pertussis* (OD_600_∼1.0) at 20,000 g for 20 min at 4°C was filtered through a syringe PES 0.22-μm filter (TPP, Switzerland) and the filtrate was precipitated overnight at 4°C by addition of trichloroacetic acid solution to a final concentration of 10 %. The precipitate was collected by centrifugation at 20,000 g for 20 min at 4°C, washed with ice-cold acetone and the dry pellet was resuspended in 50 mM ammonium bicarbonate (pH 8.3) supplemented with 8 M urea and 1 % SDS. 70 μg of total protein was applied to a 30-kDa-cutoff ultrafiltration unit, washed twice with 50 mM ammonium bicarbonate (pH 8.3) and 8 M urea followed by washing with 50 mM ammonium bicarbonate (pH 8.3) alone. Disulfide bonds were reduced by treatment of the membranes with 100 mM dithiothreitol (DTT) in 50 mM ammonium bicarbonate (pH 8.3) for 30 min at 60 °C. Subsequently, the sulfhydryl groups were alkylated by adding chloroacetamide to a final concentration of 50 mM and incubating in the dark at room temperature for 30 min. The proteins were then digested by incubation of the membranes with a MS-grade trypsin (Promega, USA) in 50 mM ammonium bicarbonate (pH 8.3) at a protein/enzyme ratio of 35:1 overnight at 37 °C. Resulting peptides were eluted from the membrane by 50 mM ammonium bicarbonate (pH 8.3) using three consecutive centrifugation steps (14,000 g for 20 min) and acidified by trifluoroacetic acid (TFA) to a final concentration of 0.1 %. The peptides were then desalted using C18 extraction disks (Empore, USA) and dried using a rotary vacuum concentrator. For LC-MS/MS analysis, the dried samples were resuspended in 2 % acetonitrile and 0.1 % TFA and loaded on an Agilent 1200 liquid chromatography system (Agilent Technologies, USA) connected to timsToF Pro mass spectrometer (Bruker, USA). The peptides (5 µl) were first trapped on C18 column (UHPLC Fully Porous Polar C18 0.3 x 20 mm, Phenomenex, USA) at a flow rate of 20 µl/min and then separated on a C18 column (Luna Omega 3 µm Polar C18 100 Å, 150 x 0.3 mm, Phenomenex, USA) using a linear gradient of acetonitrile (5-80 %) in water containing 0.1 % TFA for 35 min at a flow rate of 4 µl/min. Parameters from the standard proteomics PASEF method was used and the target intensity per individual PASEF precursor was set to 6000. The intensity threshold was set to 1500. The scan range was set between 0.6 and 1.6 V s/cm^2^ with a ramp time of 100 ms. The number of PASEF MS/MS scans was 10. Precursor ions in the *m/z* range between 100 and 1700 with charge states ≥2+ and ≤6+ were selected for fragmentation. The active exclusion was enabled for 0.4 min. Raw data were processed using PeaksStudio 10.0 software (Bioinformatics Solutions, Canada). The search parameters were set as follows: enzyme – trypsin (semispecific), oxidation of methionine and carbamidomethylation as variable modifications. The data were searched against the reference *B. pertussis* database (Uniprot: Tohama I / ATCC BAA-589 / NCTC 13251). For ‘in-gel digestion’ approach, the proteins were separated on 15 % SDS/PAGE gel and stained with Coomassie brilliant blue R-250. Protein band was excised from the gel, cut into small pieces and destained by sonication in 200 μl of Tris/HCl (pH 8.2) and 200 μl of acetonitrile (ACN). After complete destaining, the gel pieces were rinsed with 200 μl of ACN for 5 min, the liquid was discarded, and gel pieces were washed with 200 μl of distilled water. Finally, the gel was washed by 200 μl of H_2_O/ACN (1:1) and dried under vacuum. Next, the gel was rehydrated in 50 μl of 10 % ACN in 25 mM N-ethyl morpholine acetate buffer (pH 8.2) containing 1 μl of mass spectrometry-grade trypsin and let at 37 °C for 16 h. After digestion, the peptides were extracted with 100 μl of 80 % ACN, 0.1 % trifluoroacetic acid (TFA), dried via vacuum centrifugation, and solubilized in 100 μl of 2 % ACN and 0.1% formic acid. For LC-MS/MS analyses, a capillary HPLC system (1200, Agilent Technologies, Germany) connected to an ESI source of the FT-ICR mass spectrometer (15T, SolariX XR, Bruker) was used. Peptides were separated on analytical reverse phase column (MAGIC C18 AQ, 0.2×150 mm, Michrom Bioresources) and separated by following gradient: 1-10 % *B* in 1 min, 10-40 % *B* in 50 min, where solvent A was 0.2 % formic acid, 2.5 % ACN, and 2.5 % isopropanol, and solvent B was 0.16 % formic acid in 90 % ACN and 5 % isopropanol. The flow rate was 4 μl/min. ESI-FT-ICR MS was calibrated externally using arginine clusters resulting in a mass accuracy below 2 ppm. Instrument was operated in data-dependent mode, where each MS scan was followed by up to five MS/MS collision-induced fragmentations of the most intense ions. Data processing was performed using Data Analysis 4.1 (Bruker Daltonics). Peak picking was carried out by FTMS and SNAP algorithms and resulting mascot generic file was searched using local MASCOT server (MatrixScience) against the whole Swiss-Prot database. Peptide tolerance was set to 10 ppm and fragment ion tolerance to 0.05 Da and specificity for trypsin.

### SDS-PAGE and Western blotting

SDS-polyacrylamide gel electrophoresis (SDS-PAGE) was performed according to standard protocols. For Western blotting, the separated proteins were transferred to a NC 45 nitrocellulose membrane (Serva, Germany) using a Hoefer SemiPhor semi-dry transfer unit (Amersham, USA). Membranes were blocked with 5 % nonfat milk in PBS containing 0.05% Tween-20 (PBST) for 1 h at room temperature and incubated with primary antibodies in PBST overnight at 4 °C. The chemiluminiscent signal was developed on G:Box Chemi XRQ gel doc system (Syngene, USA) using a SuperSignal West Femto substrate (Thermo Fisher Scientific, USA) after incubation of the membrane with a horseradish peroxidase-conjugated goat anti-mouse or anti-rabbit secondary antibodies (Cytiva, USA) at 1:5,000 dilution in PBST for 1 h at room temperature.

## Data availability

NMR chemical shifts and distance restraints have been deposited in the Biological Magnetic Resonance Bank (BMRB) under the BMRB ID: 52089. Structural coordinates for the CT have been deposited in the Protein Data Bank under accession codes PDB ID: 8QFA. The mass spectrometry data have been deposited to the ProteomeXchange Consortium via the PRIDE partner repository with the data set identifier PXD072861. The cryo-electron tomography data are accessible through the Electron Microscopy Data Bank (EMDB) under accession code EMD-58099.

## Acknowledgement

We thank Oliva Branna and Fresia Esther Arellano Herencia for excellent technical assistance, and Attila Juhasz for support with histological analyses. This work was supported by the projects GA22-23578S, GA24-10936S and GA25-18104X of the Czech Science Foundation and AI185695 and AI191036 from the National Institutes of Health. Additional support was provided by the Ministry of Education, Youth, and Sports of the Czech Republic through the projects *Talking microbes-understanding microbial interactions within One Health framework* (CZ.02.01.01/00/22_008/0004597) and the *National Institute of Virology and Bacteriology* (LX22NPO5103). The project *Infectious diseases: new targets and strategies* within Strategy AV21 of the Czech Academy of Sciences (grant VP40) is also gratefully acknowledged. Instrumental support was provided by the research infrastructure project LM2023053 (EATRIS-CZ). We further acknowledge the Electron Microscopy Core Facility, IMG, Prague, Czech Republic, supported by projects the Ministry of Education, Youth, and Sports of the Czech Republic (LM2023050) and ERDF (CZ.02.01.01/00/23_015/0008205), and the Cryo-electron microscopy and tomography core facility CEITEC MU of CIISB, Instruct-CZ Centre, supported by projects of the Ministry of Education, Youth, and Sports of the Czech Republic (LM2023042) and ERDF (CZ.02.01.01/00/23_015/0008175).

