## Supplementary Information for "C-terminal domain of the filamentous hemagglutinin FhaB is crucial for interaction of *Bordetella pertussis* with ciliated epithelial cells"

#### Figure S1.

1 MNTNLYRLVF SHVRGMLVPV SEHCTVGNTF CGRTRGQARS GARATSLSVA PNALAWALML ACTGLPLVTH AQGLVPQGQT

81 QVLQGGNK**VP VVNIADPNSG GVSHNKFQQF NVANPGVVFN NGLTDGVSRI GGALTKNPNL TRQASAILAE VTDTSPSR**LA

161 GTLEVYGKGA DLIIANPNGI SVNGLSTLNA SNLTLTTGRP SVNGGR**IGLD VQQGTVTIER GGVNATGLGY FDVVAR**LVK**L**

241 **QGAVSSKQGK PLADIAVVAG ANRYDHATRR ATPIAAGARG AAAGAYAIDG TAAGAMYGKH ITLVSSDSGL GVR**QLGSLSS

321 PSAITVSSQG EIALGDATVQ RGPLSLK**GAG VVSAGKLASG GGAVNVAGGG AVKIASASSV GNLAVQGGGK VQATLLNAGG**

401 **TLLVSGR**QAV QLGAASSR**QA LSVNAGGALK ADK**LSATR**RV DVDGKQAVAL GSASSNALSV R**AGGALKAGK LSATGRLDVD

481 GK**QAVTLGSV ASDGALSVSA GGNLRAKQLV SSAQLEVR**GQ REVALDDASS ARGMTVVAAG ALAARNLQSK **GAIGVQGGEA**

561 **VSVANANSDA ELR**VR**GRGQV DLHDLSAAR**G ADISGEGRVN IGRARSDSDV K**VSAHGALSI DSMTALGAIG VQAGGSVSAK**

641 DMR**SRGAVTV SGGGAVNLGD VQSDGQVR**AT SAGAMTVR**DV AAAADLALQA GDALQAGFLK SAGAMTVNGR DAVRLDGAHA**

721 **GGQLRVSSDG QAALGSLAAK** GELTVSAAR**A ATVAELKSLD NISVTGGERV SVQSVNSASR VAISAHGALD VGKVSAKSGI**

801 **GLEGWGAVGA DSLGSDGAIS VSGR**DAVRVD QAR**SLADISL GAEGGATLGA VEAAGSIDVR GGSTVAANSL HANRDVR**VSG

881 K**DAVRVTAAT SGGGLHVSSG R**QLDLGAVQA R**GALALDGGA GVALQSAKAS GTLHVQGGEH LDLGTLAAVG AVDVNGTGDV**

961 **R**VAK**LVSDAG ADLQAGRSMT LGIVDTTGDL QARAQQKLEL GSVKSDGGLQ AAAGGALSLA AAEVAGALEL SGQGVTVDR**A

1041 SASRARIDST GSVGIGALK**A GAVEAASPR**R ARRALR**QDFF TPGSVVVRAQ GNVTVGRGDP HQGVLAQGDI IMDAK**GGTLL

1121 LR**NDALTENG TVTISADSAV LEHSTIESKI SQSVLAAKGD KGKPAVSVK**V AK**KLFLNGTL RAVNDNNETM SGRQIDVVDG**

1201 **RPQITDAVTG EARKDESVVS DAALVADGGP IVVEAGELVS HAGGIGNGRN KENGASVTVR TTGNLVNKGY ISAGKQGVLE**

1281 **VGGALTNEFL VGSDGTQR**IE AQRIENR**GTF QSQAPAGTAG ALVVKAAEAI VHDGVMATKG EMQIAGKGGG SPTVTAGAKA**

1361 **TTSANKLSVD VASWDNAGSL DIKKGGAQVT VAGR**YAEHGE VSIQGDYTVS ADAIALAAQV TQR**GGAANLT SRHDTRFSNK**

1441 IR**LMGPLQVN AGGAVSNTGN LK**VR**EGVTVT AASFDNETGA EVMAKSATLT TSGAARNAGK MQVKEAATIV AASVSNPGTF**

1521 **TAGK**DITVTS R**GGFDNEGKM ESNKDIVIKT EQFSNGRVLD AKHDLTVTAS GQADNRGSLK AGHDFTVQAQ RIDNSGTMAA**

1601 **GHDATLKAPH LRNTGQVVAG HDIHIINSAK LENTGR**VDAR **NDIALDVADF TNTGSLYAEH DATLTLAQGT QRDLVVDQDH**

#### Figure S1. (continued)

1681 ILPVAEGTLR VKAKSLTTEI ETGNPGSLIA EVQENIDNKQ AIVVGKDLTL SSAHGNVANE ANALLWAAGE LTVKAQNITN

1761 KRAALIEAGG NARLTAAVAL LNKLGRIRAG EDMHLDAPRI ENTAKLSGEV QRKGVQDVGG GEHGRWSGIG YVNYWLRAGN

1841 GKKAGTIAAP WYGGDLTAEQ SLIEVGKDLY LNAGARKDEH RHLLNEGVIQ AGGHGHIGGD VDNRSVVRTV SAMEYFKTPL

1921 PVSLTALDNR AGLSPATWNF QSTYELLDYL LDQNRYEYIW GLYPTYTEWS VNTLKNLDLG YQAKPAPTAP PMPKAPELDL

2001 RGHTLESAEG RKIFGEYKKL QGEYEKAKMA VQAVEAYGEA TRRVHDQLGQ RYGKALGGMD AETKEVDGII QEFAADLRTV

2081 YAKQADQATI DAETDKVAQR YKSQIDAVRL QAIQPGRVTL AKALSAALGA DWRALGHSQL MQRWKDFKAG KRGAEIAFYP

2161 KEQTVLAAGA GLTLSNGAIH NGENAAQNRG RPEGLKIGAH SATSVSGSFD ALRDVGLEKR LDIDDALAAV LVNPHIFTRI

2241 GAAQTSLADG AAGPALARQA RQAPETDGMV DARGLGSADA LASLASLDAA QGLEVSGRRN AQVADAGLAG PSAVAAPAVG

2321 AADVGVEPVT GDQVDQPVVA VGLEQPVATV RVAPPAVALP RPLFETRIKF IDQSKFYGSR YFFEQIGYKP DRAARVAGDN

2401 YFDTTLVREQ VRRALGGYES RLPVRGVALV AKLMDSAGTV GKALGLKVGV APTAQQLKQA DRDFVWYVDT VIDGQKVLAP

2481 RLYLTEATRQ GITDQYAGGG ALIASGGDVT VNTDGHDVSS VNGLIQGRSV KVDAGKGKVV VADSKGAGGG IEADDEVDVS

2561 GRDIGIEGGK LRGKDVRLKA DTVKVATSMR YDDKGRLAAR GDGALDAQGG QLHIEAKRLE TAGATLKGGK VKLDVDDVKL

2641 GGVYEAGSSY ENKSSTPLGS LFAILSSTTE TNQSAHANHY GTRIEAGTLE GKMQNLEIEG GSVDAAHTDL SVARDARFKA

2721 AADFAHAEHE KDVRQLSLGA KVGAGGYEAG FSLGSESGLE AHAGRGMTAG AEVKVGYRAS HEQSSETEKS YRNANLNFGG

2801 GSVEAGNVLD IGGADINRNR YGGAAKGNAG TEEALRMRAK KVESTKYVSE QTSQSSGWSV EVASTASARS SLLTAATRLG

2881 DSVAQNVEDG REIRGELMAA QVAAEATQLV TADTAAVALS AGISADFDSS HSRSTSQNTQ YLGGNLSIEA TEGDATLVGA

2961 KFGGGDQVSL KAAKSVNLMA AESTFESYSE SHNFHASADA NLGANAVQGA VGLGLTAGMG TSHQITNETG KTYAGTSVDA

3041 ANVSIDAGKD LNLSGSRVRG KHVVLDVEGD INATSKQDER NYNSSGGGWD ASAGVAIQNR TLVAPVGSAG FNFNTEHDNS

3121 RLTNDGAAGV VASDGLTGHV KGDANLTGAT IADLSGKGNL KVDGAVNAQN LKDYRDKDGG SGGLNVGISS TTLAPTVGVA

3201 FGRVAGEDYQ AEQRATIDVG QTKDPARLQV GGGVKGTLNQ DAAQATVVQR NKHWAGGGSE FSVAGKSLKK KNQVRPVETP

3281 TPDVVDGPPS RPTTPPASPQ PIRATVEVSS PPPVSVATVE VVPRPKVETA QPLPPRPVAA QVVPVTPPKV EVAKVEVVPR

3361 PKVETAQPLP PRPVVAEKVT TPAVQPQLAK VETVQPVKPE TTKPLPKPLP VAKVTKAPPP VVETAQPLPP VKPQKATPGP

3441 VAEVGKATVT TVQVQSAPPK PAPVAKQPAP APKPKPKPKP KAERPKPGKT TPLSGRHVVQ QQVQVLQRQA SDINNTKSLP

3521 GGKLPKPVTV KLTDENGKPQ TYTINRREDL MKLNGKVLST KTTLGLEQTF RLRVEDIGGK NYRVFYETNK

**Figure S1. Tryptic peptide coverage map of FhaB detected in the bacterial lysate of *B. pertussis* grown in liquid culture.** Bacterial cells were collected by centrifugation of *B. pertussis* culture grown in SS medium to $OD_{600\ nm}$ of 1.0. Proteins were digested with trypsin, and the resulting peptides were analyzed by mass spectrometry. Detected peptides are indicated by blue lines, and the corresponding covered regions of the FhaB sequence are highlighted in grey. Oxidized methionine residues are shown in red.

#### Figure S2.

1 MNTNLYRLVF SHVRGMLVPV SEHCTVGNTF CGRTRGQARS GARATSLSVA PNALAWALML ACTGLPLVTH AQGLVPQGQT

81 QVLQGGNKVP VVNIADPNSG GVSHNKFQQF NVANPGVVFN NGLTDGVSRI GGALTKNPNL TRQASAILAE VTDTSPSRLA

161 GTLEVYGKGA DLIIANPNGI SVNGLSTLNA SNLTLTTGRP SVNGGRIGLD VQQGTVTIER GGVNATGLGY FDVVARLVKL

296

241 QGAVSSKQGK PLADIAVVAG ANRYDHATRR ATPIAAGARG AAAGAYAIDG TAAGAMYGKH ITLVSSDSGL GVRQLGSLSS

321 PSAITVSSQG EIALGDATVQ RGPLSLKGAG VVSAGKLASG GGAVNVAGGG AVKIASASSV GNLAVQGGGK VQATLLNAGG

401 TLLVSGRQAV QLGAASSRQA LSVNAGGALK ADKLSATRRV DVDGKQAVAL GSASSNALSV RAGGALKAGK LSATGRLDVD

481 GKQAVTLGSV ASDGALSVSA GGNLRAKQLV SSAQLEVRGQ REVALDDASS ARGMTVVAAG ALAARNLQSK GAIGVQGGEA

561 VSVANANSDA ELRVRGRGQV DLHDLSAARG ADISGEGRVN IGRARSDSDV KVSAHGALSI DSMTALGAIG VQAGGSVSAK

675 705

641 DMRSRGAVTV SGGGAVNLGD VQSDGQVRAT SAGAMTVRDV AAAADLALQA GDALQAGFLK SAGAMTVNGR DAVRLDGAHA

721 GGQLRVSSDG QAALGSLAAK GELTVSAARA ATVAELKSLD NISVTGGERV SVQSVNSASR VAISAHGALD VGKVSAKSGI

801 GLEGWGAVGA DSLGSDGAIS VSGRDAVRVD QARSLADISL GAEGGATLGA VEAAGSIDVR GGSTVAANSL HANRDVRVSG

881 KDAVRVTAAT SGGGLHVSSG RQLDLGAVQA RGALALDGGA GVALQSAKAS GTLHVQGGEH LDLGTLAAVG AVDVNGTGDV

979

961 RVAKLVSDAG ADLQAGRSMT LGIVDTTGDL QARAQQKLEL GSVKSDGGLQ AAAGGALSLA AAEVAGALEL SGQGVTVDRA

1041 SASRARIDST GSVGIGALKA GAVEAASPRR ARRALRQDFF TPGSVVVRAQ GNVTVGRGDP HQGVLAQGDI IMDAKGGTLL

1190

1121 LRNDALTENG TVTISADSAV LEHSTIESKI SQSVLAAKGD KGKPAVSVKV AKKLFLNGTL RAVNDNNETM SGRQIDVVDG

1201 RPQITDAVTG EARKDESVVS DAALVADGGP IVVEAGELVS HAGGIGNGRN KENGASVTVR TTGNLVNKGY ISAGKQGVLE

1336 1342

1281 VGGALTNEFL VGSDGTQRIE AQRIENRGTF QSQAPAGTAG ALVVKAAEAI VHDGVMATKG EMQIAGKGGG SPTVTAGAKA

1361 TTSANKLSVD VASWDNAGSL DIKKGGAQVT VAGRYAEHGE VSIQGDYTVS ADAIALAAQV TQRGGAANLT SRHDTRFSNK

1444

1441 IRLMGPLQVN AGGAVSNTGN LKVREGVTVT AASFDNETGA EVMAKSATLT TSGAARNAGK MQVKEAATIV AASVSNPGTF

1598

1521 TAGKDITVTS RGGFDNEGKM ESNKDIVIKT EQFSNGRVLD AKHDLTVTAS GQADNRGSLK AGHDFTVQAQ RIDNSGTMAA

1601 GHDATLKAPH LRNTGQVVAG HDIHIINSAK LENTGRVDAR NDIALDVADF TNTGSLYAEH DATLTLAQGT QRDLVVDQDH

1681 ILPVAEGTLR VKAKSLTTEI ETGNPGSLIA EVQENIDNKQ AIVVGKDLTL SSAHGNVANE ANALLWAAGE LTVKAQNITN

1793

1761 KRAALIEAGG NARLTAAVAL LNKLGRIRAG EDMHLDAPRI ENTAKLSGEV QRKGVQDVGG GEHGRWSGIG YVNYWLRAGN

#### Figure S2. (continued)

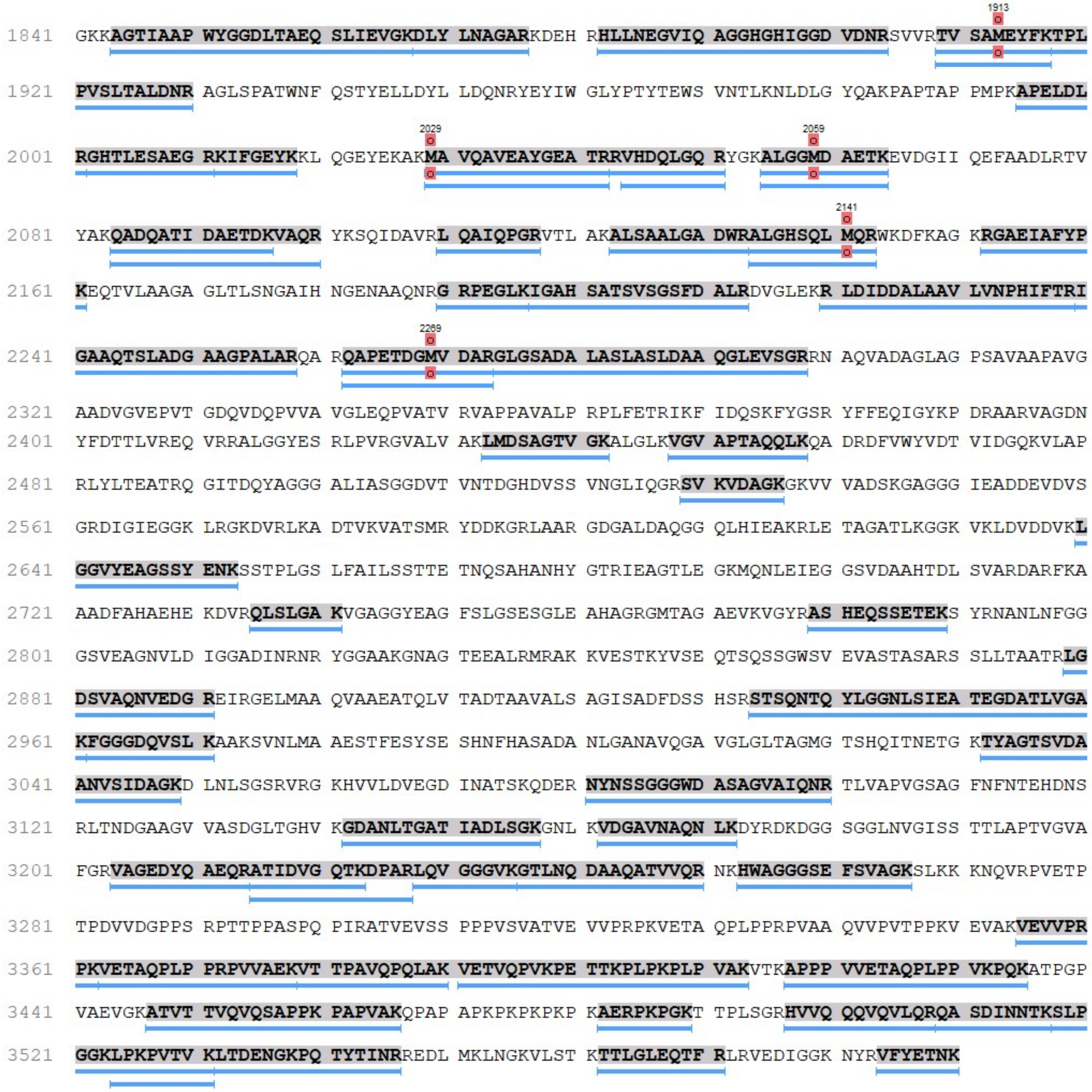


**Figure S2. Tryptic peptide coverage map of FhaB detected in the culture supernatant of *B. pertussis* grown in liquid culture.** Culture supernatants were collected by centrifugation of *B. pertussis* culture grown in SS medium to $OD_{600\ nm}$ of 1.0. Proteins were precipitated with trichloroacetic acid, digested with trypsin, and the resulting peptides were analyzed by mass spectrometry. Detected peptides are indicated by blue lines, and the corresponding covered regions of the FhaB sequence are highlighted in grey. Oxidized methionine residues are shown in red.

**Figure S3.**

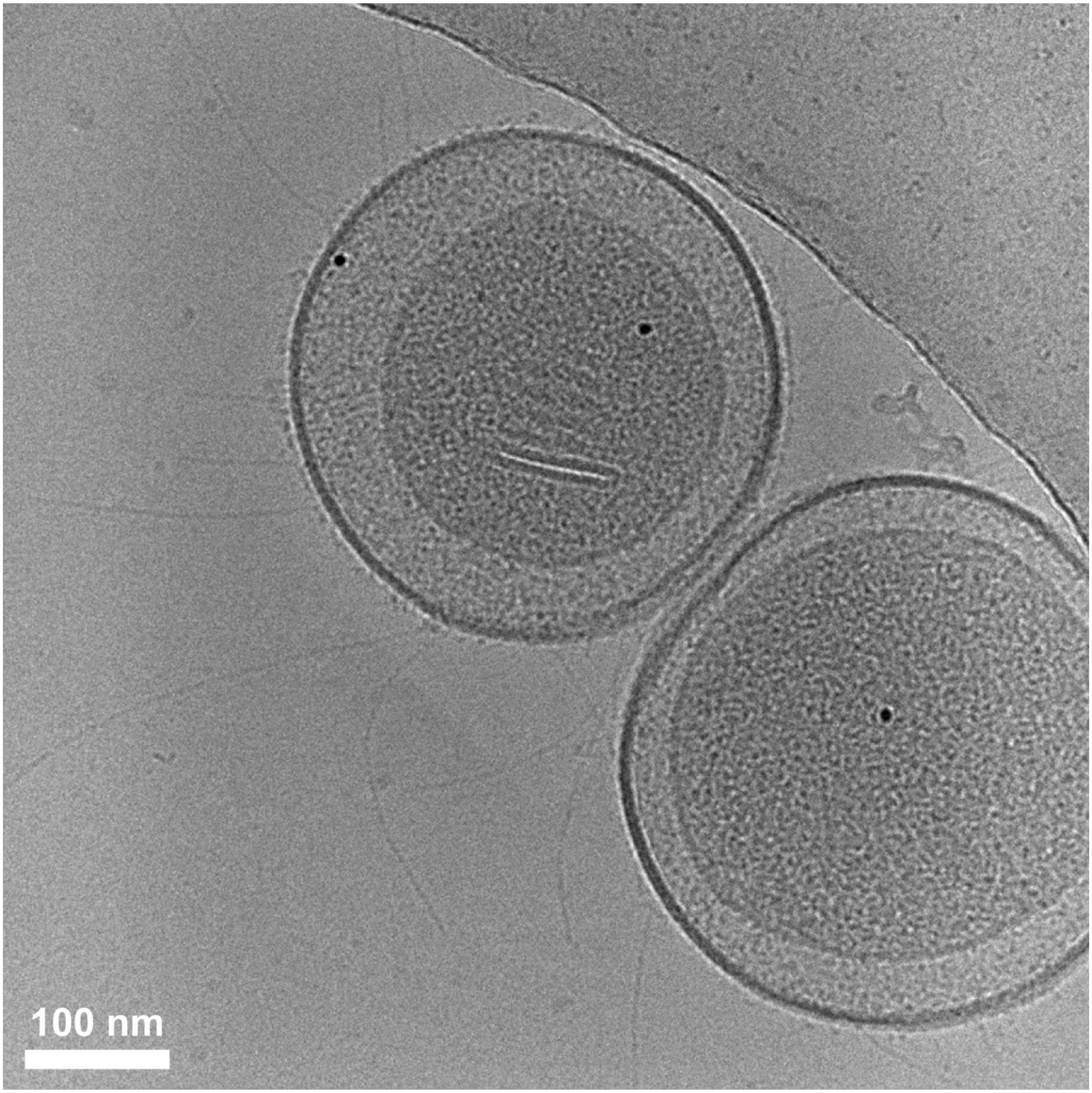


**Figure S3. Cryo-EM image of minicells derived from the wild-type *B. pertussis* strain.** Minicells were purified by a combination of differential and density gradient centrifugation following overexpression of the *ftsZAQ* genes in *B. pertussis* grown in liquid culture at 37 °C, and imaged under cryogenic conditions using a Jeol JEM-F200 transmission electron microscope equipped with TemCam XF416 camera. Black dots correspond to 5-nm gold nanoparticles.

**Figure S4.**

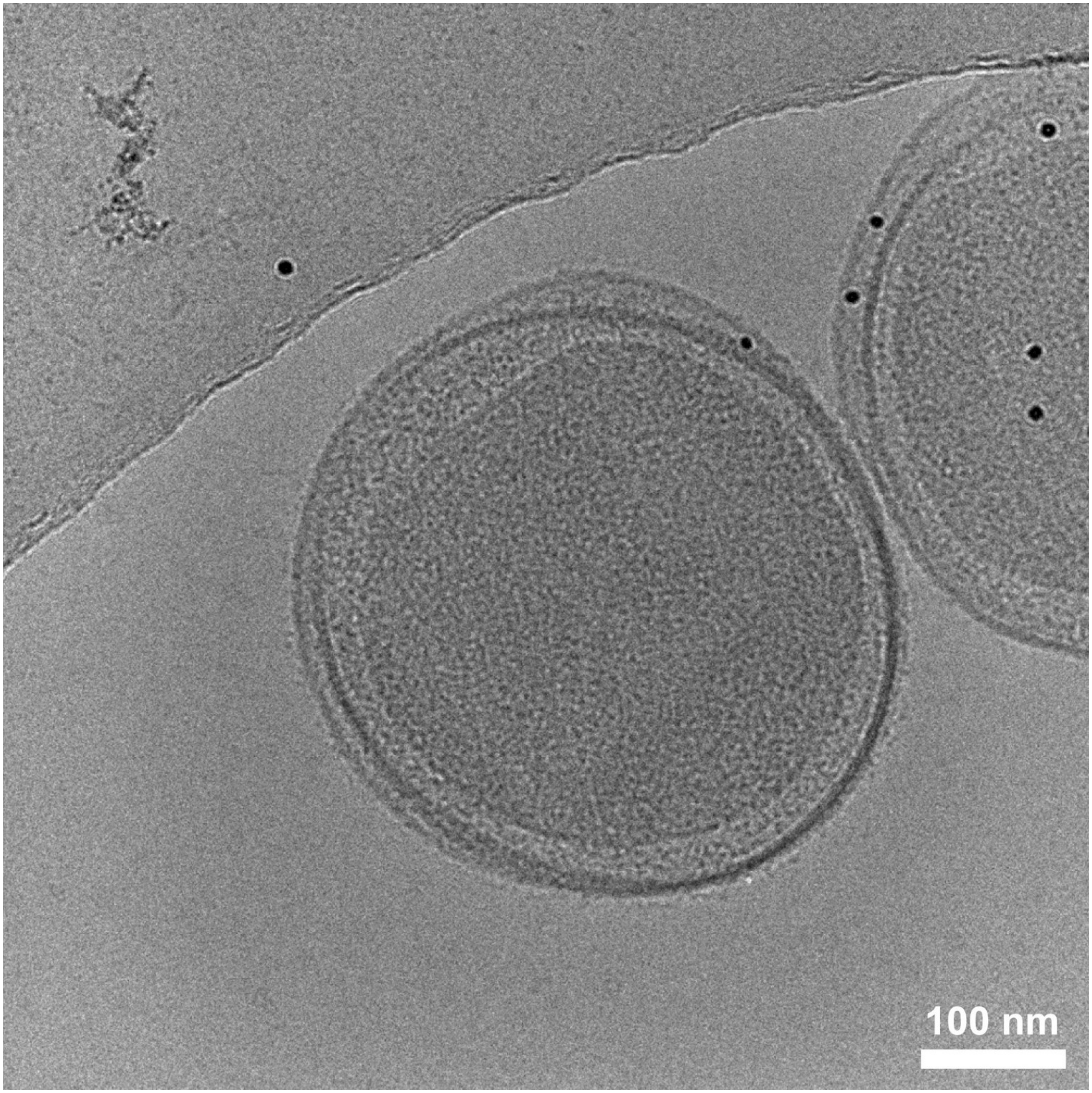


**Figure S4. Cryo-EM image of minicells derived from the Δ*fimA-D* strain of *B. pertussis.*** Minicells were purified by a combination of differential and density gradient centrifugation following overexpression of the *ftsZAQ* genes in the Δ*fimA-D/sphB1*⁺ *B. pertussis* strain grown in liquid culture at 37 °C, and imaged under cryogenic conditions using a Jeol JEM-F200 transmission electron microscope equipped with TemCam XF416 camera. Proteolytic activity of SphB1 inhibits the persistence of FhaB filaments on the cell suface. Black dots correspond to 5-nm gold nanoparticles.

**Figure S5.**

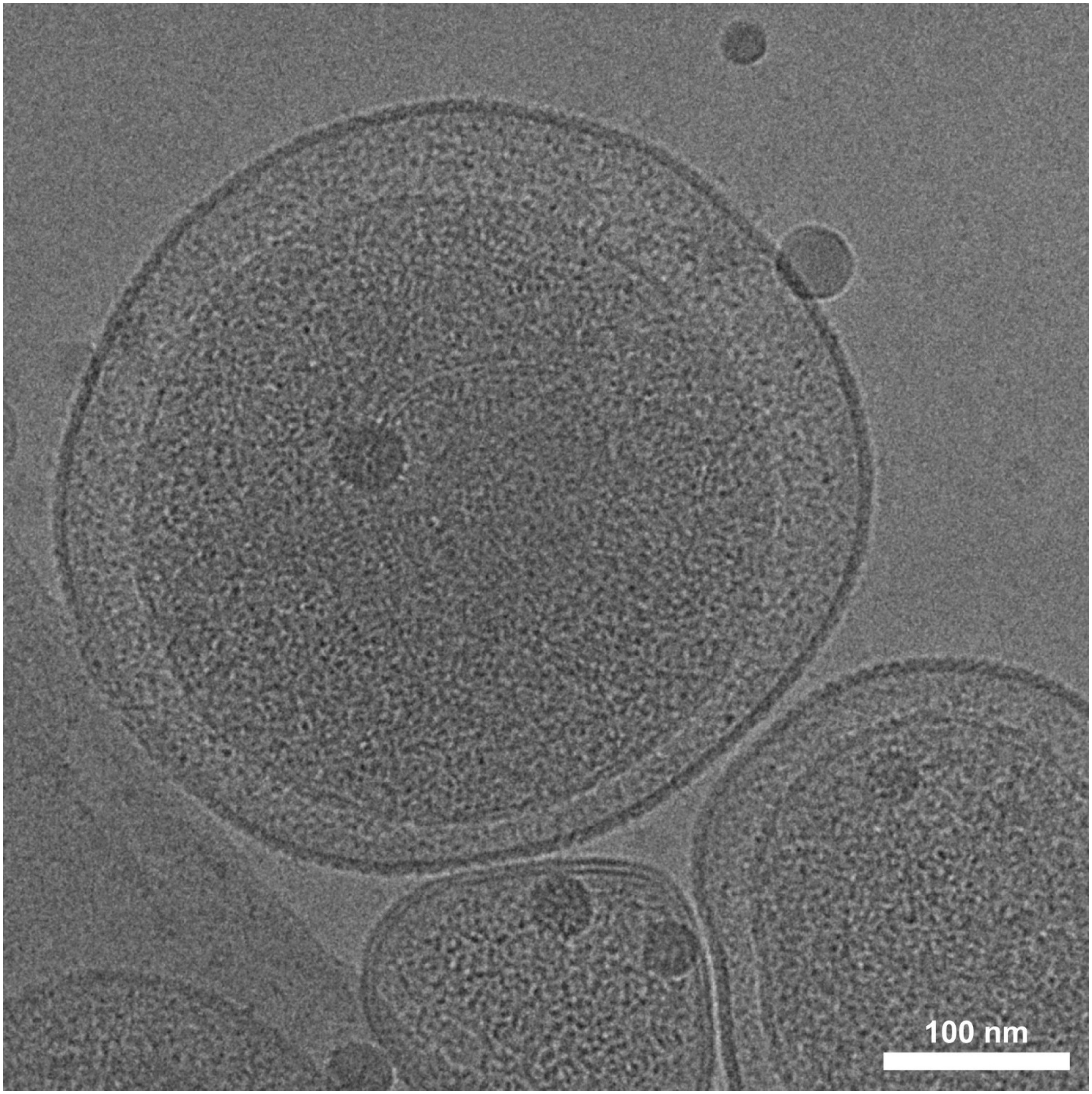


**Figure S5. Cryo-EM image of minicells derived from the Δ*fhaB* strain of *B. pertussis.*** Minicells were purified by a combination of differential and density gradient centrifugation following overexpression of the *ftsZAQ* genes in the *fhaB*-deficient strains of *B. pertussis* carrying a Δ*sphB1*/Δ*fimA-D* background, and imaged under cryogenic conditions using a Talos Arctica transmission electron microscope equipped with a Falcon 4 direct electron detector.

### Figure S6.

**A**

```
                           ********************************************************
      B. pertussis 3495 GRHVVQQQVQVLQRQASDINNTKSLPGGKLPKPVTVKLTDENGKPQTYTINRREDL 3550
B. bronchiseptica 3615 GRHVVQQQVQVLQRQASDINNTKSLPGGKLPKPVTVKLTDENGKPQTYTINRREDL 3670

                           ****************************************
      B. pertussis 3551 MKLNGKVLSTKTTLGLEQTFRLRVEDIGGKNYRVFYETNK 3590
B. bronchiseptica 3671 MKLNGKVLSTKTTLGLEQTFRLRVEDIGGKNYRVFYETNK 3710
```

**B**

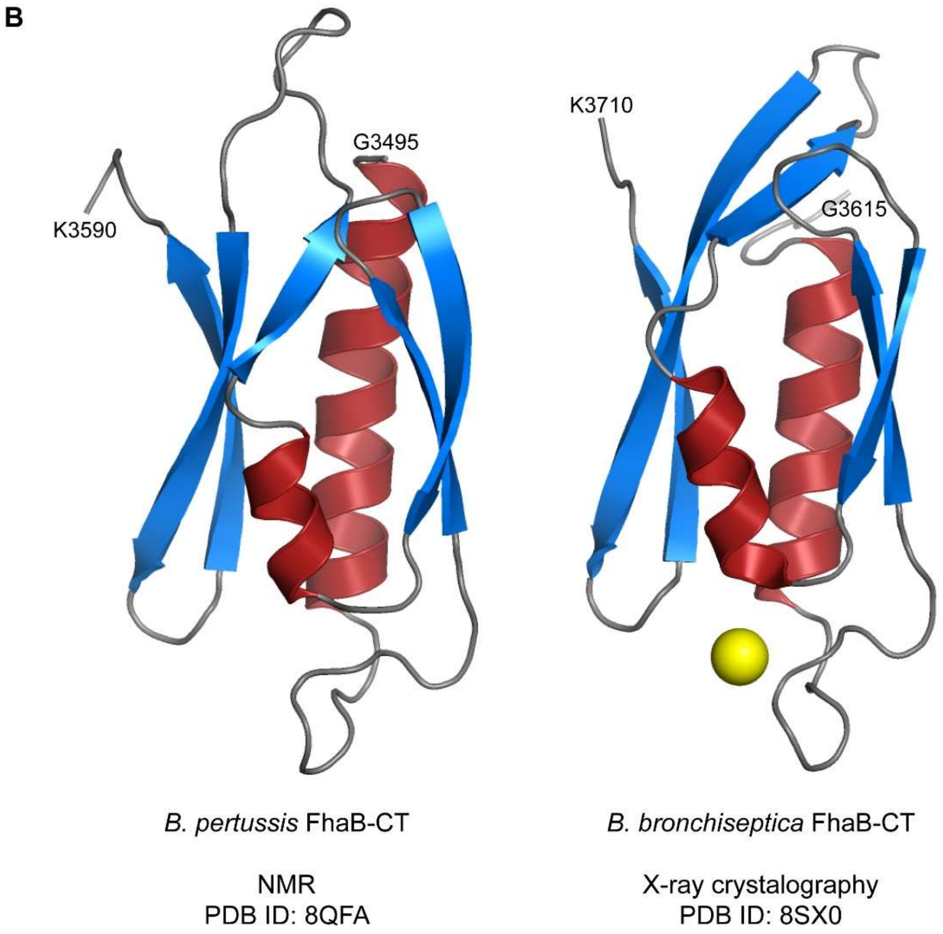


**Figure S6. Sequence and structural comparison of the C-terminal domains of FhaB from *B. pertussis* and *B. bronchiseptica*.** (**A**) Alignment of the primary amino acid sequences of the C-terminal domains of FhaB from *B. pertussis* Tohama I (NCBI: WP_010930610.1) and *B. bronchiseptica* RB50 (WP_041936401.1), generated using Clustal Omega (https://www.ebi.ac.uk/jdispatcher/msa/clustalo). Asterisks (*) above the alignment indicate conserved residues. (**B**) Structural comparison of the C-terminal domains of *B. pertussis* and *B. bronchiseptica*, determined by NMR spectroscopy and X-ray crystallography, respectively. The structures are shown in cartoon representation, with α-helices colored red and β strands colored blue. Yellow sphere represents $Ca^{2+}$ ion.

**Figure S7.**

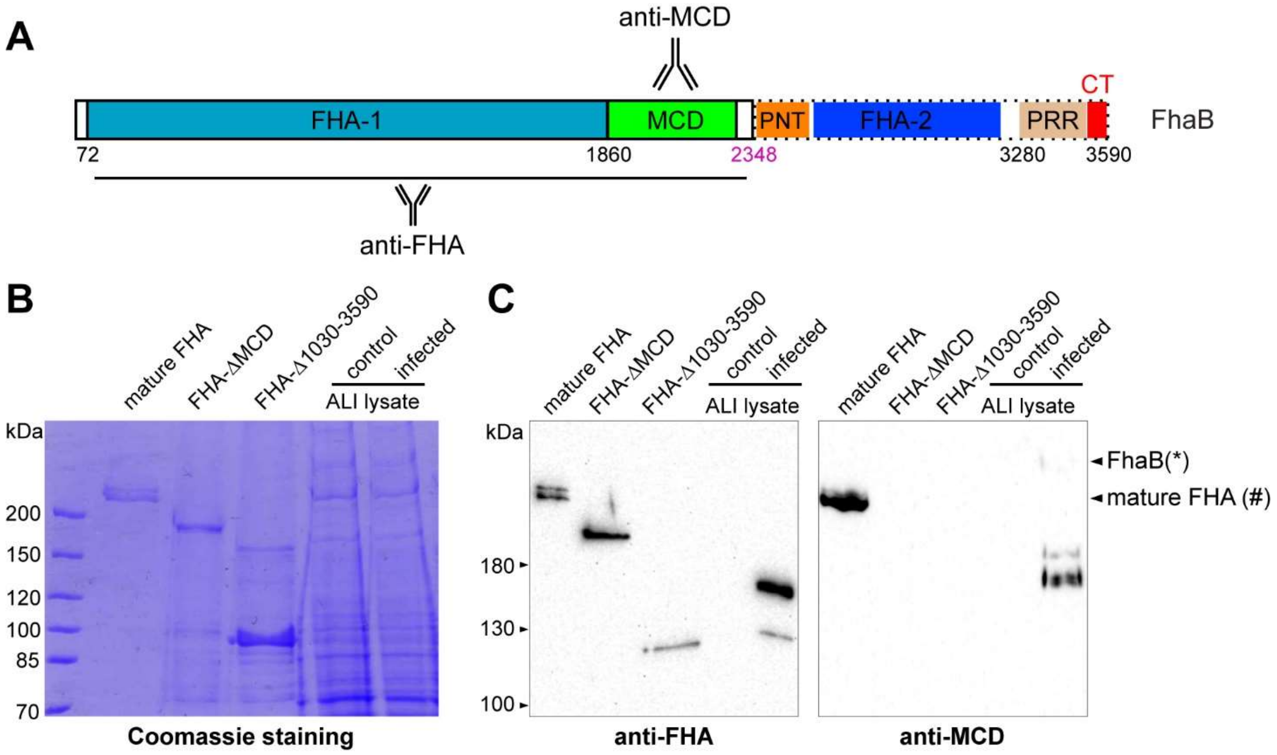


**Figure S7. FhaB is processed at its N terminus following interaction of *B. pertussis* with ciliated epithelial cells.** (**A**) Epitope map of polyclonal anti-FHA and anti-MCD sera on FhaB. FHA-1 and FHA-2 denote regions containing tandem 19–amino acid repeats predicted to form a right-handed β-helical shaft. MCD, mature C-terminal domain; PNT, prodomain N terminus; CT, C-terminal domain. (**B**) Coomasie-stained SDS-PAGE gel (5 %) of purified mature FHA, FHA-ΔMCD, and FHA-Δ1030-3590 proteins as well as lysates from control and *B. pertussis*-infected ALI-differentiated airway epithelial cells. FHA proteins were purified from culture supernatants of the indicated *B. pertussis* strains by affinity chromatography using Cellufine Sulfate. ALI-differentiated airway epithelial cells were infected with *B. pertussis* ($5 \times 10^5$ CFU in 50 µl HBSS per transwell) for 16 hours and then maintained for an additional 24 h under ALI conditions. (**C**) Immunoblot analyses of samples shown in (B), probed with polyclonal anti-FHA and anti-MCD serum.
